# Model-based hearing-enhancement strategies for cochlear synaptopathy pathologies

**DOI:** 10.1101/2022.01.10.475652

**Authors:** Fotios Drakopoulos, Viacheslav Vasilkov, Alejandro Osses Vecchi, Tijmen Wartenberg, Sarah Verhulst

**Author notes:** Department of Otolaryngology – Head & Neck Surgery, Harvard Medical School, Massachusetts Eye & Ear, Boston, Massachusetts, USA. Laboratoire des systèmes perceptifs, Département d’études cognitives, École Normale Supérieure (ENS), PSL University, Paris, France. Move the Brain, Zeist, Utrecht, the Netherlands (Tijmen Wartenberg).

## Abstract

It is well known that ageing and noise exposure are important causes of sensorineural hearing loss, and can result in damage of the outer hair cells or other structures of the inner ear, including synaptic damage to the auditory nerve (AN), i.e., cochlear synaptopathy (CS). Despite the suspected high prevalence of CS among people with self-reported hearing difficulties but seemingly normal hearing, conventional hearing-aid algorithms do not compensate for the functional deficits associated with CS. Here, we present and evaluate a number of auditory signal-processing strategies designed to maximally restore AN coding for listeners with CS pathologies. We evaluated our algorithms in subjects with and without suspected age-related CS to assess whether physiological and behavioural markers associated with CS can be improved. Our data show that after applying our algorithms, envelope-following responses and perceptual amplitude-modulation sensitivity were consistently enhanced in both young and older listeners. Speech-in-noise intelligibility showed small improvements after processing but mostly for young normal-hearing participants, with median improvements of up to 8.3%. Since our hearing-enhancement strategies were designed to optimally drive the AN fibres, they were able to improve temporal-envelope processing for listeners both with and without suspected CS. Our proposed algorithms can be rapidly executed and can thus extend the application range of current hearing aids and hearables, while leaving sound amplification unaffected.

## 1 Introduction

With age, our hearing ability starts to decline: Communicating in noisy environments becomes challenging, and the hearing of faint sounds difficult. Even though there has been extensive research on the understanding and restoration of hearing [1, 2], a precise diagnostic assay of the various aspects that constitute hearing impairment is still unavailable. The resulting mixed success in the treatment of sensorineural hearing loss (SNHL) reveals an incomplete understanding of how each component of the auditory system affects human sound perception. This was recently demonstrated by the ground-breaking discovery of cochlear synaptopathy (CS) [3–5], which showed that ageing [6, 7] and noise exposure [8] can cause a selective and substantial loss of auditory nerve fibres (ANFs) in animals as well as humans. Evidence from human and animal studies suggests that CS impacts the encoding of speech in everyday listening conditions and the comprehension of speech in noise [9–11]. The functional consequences of CS have also been referred to as “hidden hearing loss” [12], as CS cannot be detected using standard audiometric threshold measures. In animal models, the onset of CS occurs earlier in time than that of the outer-hair-cell (OHC) loss that is associated with elevated hearing thresholds [3, 4], suggesting that a large portion of the noise-exposed and ageing population may suffer from undiagnosed CS, while their audibility falls within the normal audiometric range.

Recent studies have shown that CS mainly affects the robust processing of the temporal-envelope information in sound [13,14]. Functional consequences of synaptopathy include a reduced neural representation of instantaneous supra-threshold amplitude fluctuations [10], and stochastic under-sampling of speech in noisy acoustic scenarios [15, 16]. Despite the suspected high prevalence of CS in listeners with self-reported hearing difficulties, or those with OHC damage, conventional hearing-aid algorithms do not specifically compensate for the functional deficits associated with CS. On the contrary, hearing aids usually rely on non-linear dynamic-range compression algorithms that reduce amplitude fluctuations of the temporal envelope [17–19], possibly resulting in a further deterioration of hearing perception in individuals with CS [20–22]. Thus, a large population of those with self-reported hearing difficulties but normal audiograms, or even those with impaired audiograms, may benefit from auditory signal-processing algorithms that specifically target the functional deficits associated with CS.

Since temporal-envelope coding is degraded by CS, a prime candidate for acoustic intervention is thus to modify the speech envelope which is essential for robust speech intelligibility (SI) [23–31]. Acoustic modifications of the speech-envelope shape and periodicity can improve SI [18,32–37], especially in connection with hearing through cochlear implants [38–43]. Several studies have shown that envelope enhancement can be beneficial for speech recognition in individuals with temporal processing deficits [22, 36, 44–48]. Here, we go beyond these conventional approaches to develop a model-based type of auditory envelope processing that operates directly on the signal waveform to counteract the functional consequences associated with CS. We focus on CS-compensating signal-processing algorithms that enhance the temporal envelope of the signal while avoiding effects on the temporal fine-structure information, to avoid possible decrements in SI [49].

To develop this new type of CS-compensating audio processing, we adopted a biophysically inspired computational model of the human auditory periphery [50] that simulates how CS impacts auditory sound processing. Even though individualised models of hearing-impaired auditory processing are widely adopted in the design of hearing-aid signal processing [51–54], they are typically based on the compensation of degraded hearing thresholds or of perceived loudness, to yield optimal acoustic amplification to hearing-aid users [55, 56]. Instead, we devised a computational optimisation method to design a number of envelope-processing algorithms that maximally restored simulated CS-affected auditory-nerve (AN) responses. We evaluated our model-designed algorithms in participants with and without suspected age-related CS, to test whether our algorithms enhanced a number of biophysical and perceptual markers associated with CS. These markers included envelope-following responses (EFRs), amplitude-modulation (AM) detection sensitivity, and SI in terms of speech reception thresholds (SRTs) and word recognitions scores (WRSs). Because a direct quantification of CS relies on post-mortem imaging of the temporal bone [7, 57], we focussed on older listeners who had been shown to be affected by CS through non-invasive electroencephalographic (EEG) techniques [14, 58, 59].

## 2 Design of the hearing-enhancement algorithms

We developed our hearing-enhancement algorithms based on a biophysically inspired auditory periphery model [50] that can simulate human auditory processing in normal-hearing (NH) and hearing-impaired (HI) subjects. The peripheral auditory model includes a transmission-line description of the cochlea and an inner-hair-cell/auditory-nerve synaptic complex model to generate AN firing rates across a number of cochlear tonotopic locations [50]. We aimed to design signal-processing strategies that can restore peripheral AN processing in CS-affected peripheries, and thereby ultimately improve speech perception and intelligibility. A general description of the stages we followed for the design of the hearing-restoration algorithms is given here and the reader is referred to Methods (Sec. 7) for a more detailed description.

### 2.1 Optimisation procedure

We applied an iterative optimisation approach (Fig. 1) to derive an auditory-processing algorithm that minimises the difference between simulated NH and CS-affected AN responses. The model-simulated AN responses corresponded to the summed responses of the ANF population present in a NH and a CS-affected periphery, respectively. The derivative-free optimisation that we applied is efficient at discovering local minima of continuous and smooth functions for constrained problems, but it cannot guarantee a reliable solution when the problem is more complex. To this end, we constrained the optimisation problem by using a simple stimulus, i.e., a sinusoidally-amplitude-modulated (SAM) pure tone with a carrier frequency *f*_*c*_ = 4 kHz, modulation frequency *f*_*m*_ = 120 Hz and *m* = 100% modulation, that can be related to many behavioural and physiological studies of temporal-envelope processing in humans [14, 60–62].

**Fig 1.**
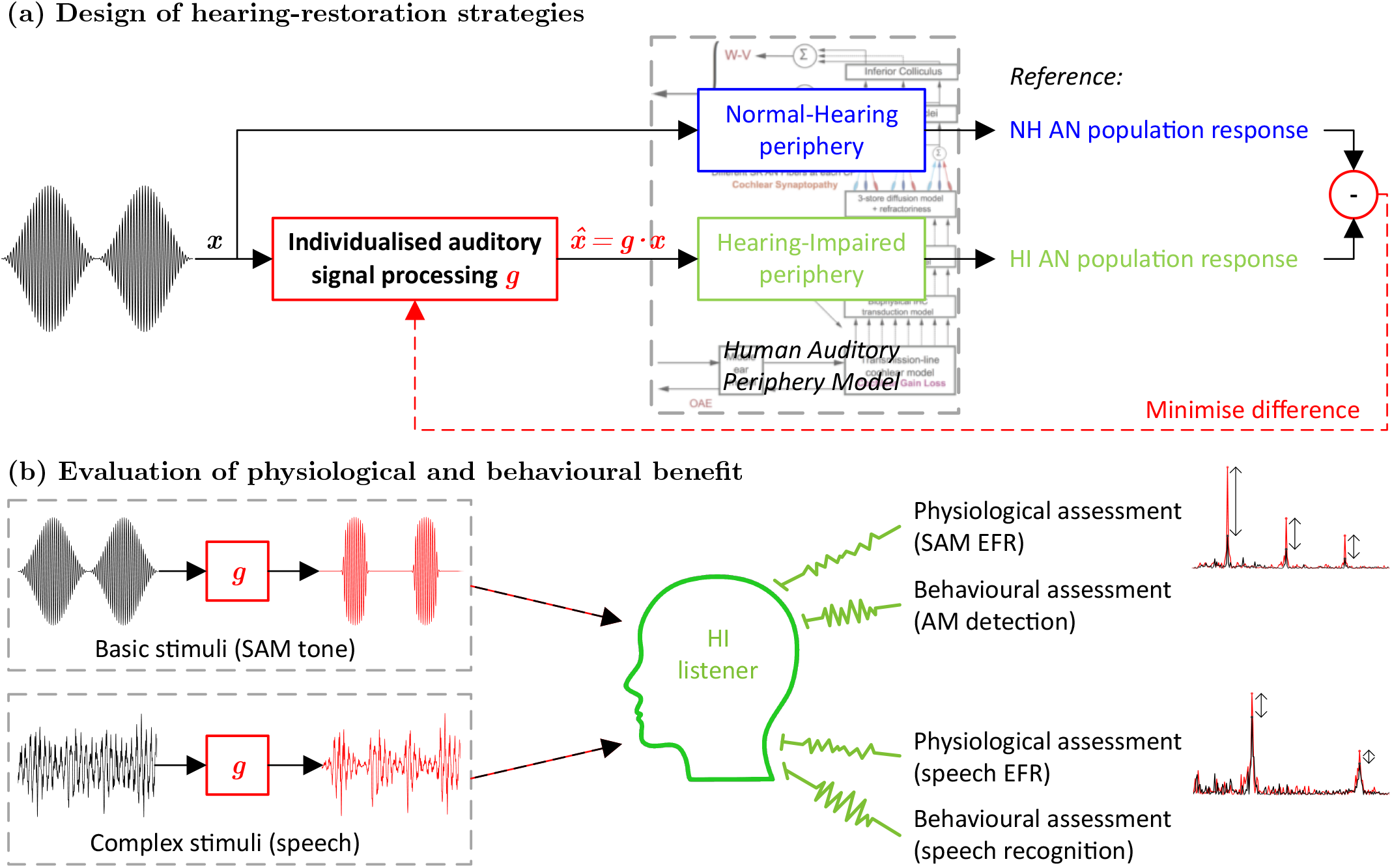
Framework for the design and evaluation of hearing-restoration strategies. **a** The schematic shows how signal-processing strategies can be designed to maximally restore a hearing-impaired (HI) AN response to the reference (NH) response. In our case, the HI periphery corresponded to a CS-affected periphery and the auditory signal processing to a non-linear envelope-processing function *g*. The parameters of the processing function *g* were optimised to generate a processed signal that minimises the difference between the two responses. The same optimisation approach can be applied to any HI periphery with different degrees of CS and/or OHC loss. **b** By applying the processing function *g* to different stimuli, the physiological and behavioural benefit of the processed sounds can be assessed on HI listeners. Ideally, the SNHL auditory profile of the HI listener corresponds to the simulated HI periphery that was used in the optimisation (**a**), yielding an individualised hearing restoration

### 2.2 CS compensation

The optimisation procedure was applied to three peripheries that included different degrees of CS, abbreviated as 13,0,0, 10,0,0 and 7,0,0, with each number corresponding to the quantity of high (H), medium (M) and low (L) spontaneous-rate (SR) ANF innervations, respectively. When compared to the reference NH periphery (13,3,3 ANFs [50, 63]), these corresponded to a complete loss of LSR and MSR ANFs, but with either healthy HSR ANFs (13,0,0), or with losses of 23% (10,0,0) or 46% (7,0,0) of the HSR ANFs, respectively. For each CS profile, an auditory signal-processing function *g* was optimised to maximally enhance the respective CS-affected AN output in response to the processed stimulus (Fig. 1a).

### 2.3 Auditory signal processing

We chose an envelope-processing function *g* (Eq. *2*) that preserves the stimulus envelope peaks, but modifies sound onsets and offsets to increase the resting periods between stimulation. The (remaining) ANFs are thus able to recover faster from prior stimulation to generate stronger onset responses and more synchronised AN activity [14, 32, 38, 39, 42, 64, 65]. We focussed on enhancing the steady-state responses to temporal peaks in the stimulus (see Methods), thus excluding the initial onset peak (and the subsequent exponential decay) of the ANF responses that was shown to be enhanced after CS in recent animal studies [66]. During each iteration of the optimisation procedure, the parameters of the signal-processing function *g* were adjusted so that it could generate a processed stimulus 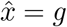 · *x* that minimised the difference between the NH and the respective CS-affected AN response. The selected non-linear function *g* was easy to implement and was executed online (*<*5 ms latency for a 50-ms stimulus in MATLAB), allowing for future extensions in real-time applications.

Figure 2a shows the three processing functions that were derived after applying the optimisation procedure for the three selected CS profiles. The processing functions are shown as a function of the amplitude of the input stimulus *x*, normalised to its maximum absolute (peak) amplitude. Thus, for each CS profile, the envelope of the unprocessed SAM stimulus of Fig. 2b (modulating tone) can be used to compute the processing function *g* across time (Eq. *2*), which is then multiplied elementwise by the input stimulus *x* to derive the processed version 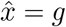 · *x*. The processed signal 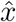 is able to optimally stimulate the corresponding CS-affected periphery to generate an enhanced AN response that closely matches the NH response. Figure 2c demonstrates that simulated CS-affected AN responses can be partially or fully restored after applying our processing to fully-modulated SAM tones.

**Fig 2.**
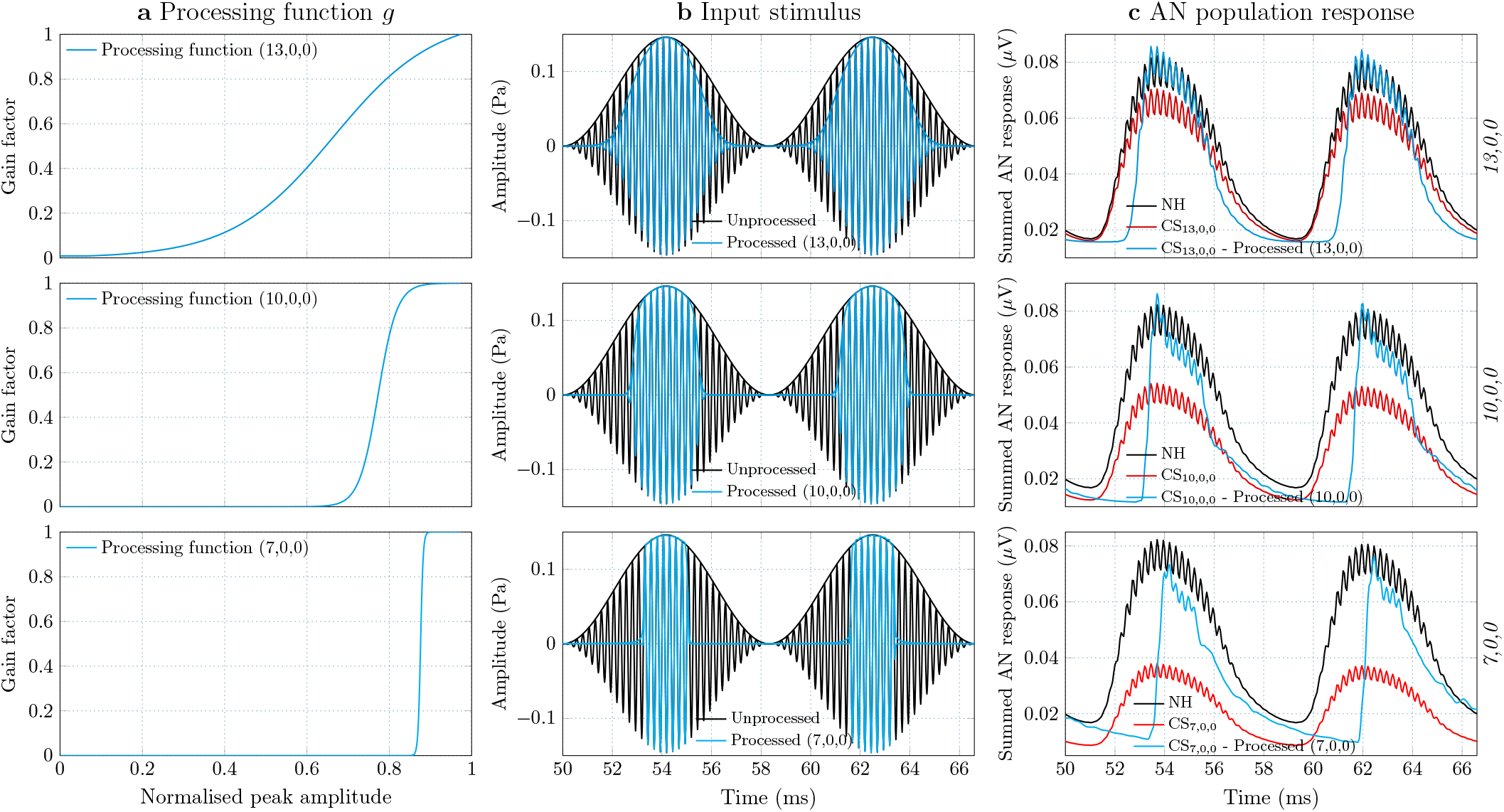
CS compensation of a fully-modulated SAM tone. Enhancement of the AN responses of three CS-affected peripheries to a fully-modulated 70-dB-SPL SAM tone stimulus. **a** Three non-linear functions were derived from our optimisation method that maximally restore the AN responses for three degrees of CS. **b** Based on the signal envelope, the non-linear functions were applied to the SAM tones across time to derive the processed stimuli for each CS type. **c** The simulated AN responses of the NH and CS peripheries are shown in response to the unprocessed SAM stimulus (black and red) and the respective processed version (cyan), when given as input to the corresponding CS periphery.

### 2.4 Generalising the hearing-enhancement algorithms

Even though the algorithms were designed to restore ANF coding to fully-modulated SAM tones, the same processing can be applied to stimuli with different degrees of modulation, or to speech, which naturally contains envelope fluctuations of different strength. However, when applying the envelope-processing functions of Fig. 2a to partially-modulated stimuli, the modulation depth of the stimulus envelope will also be increased. This is not always desired, especially when the aim is to compare the effect of ANF stimulation recovery on the responses without affecting the dynamic range. Hence, we also designed a modified strategy *g*_*m*_ that only processes the slope of the modulation envelope while preserving the modulation depth of the original stimulus intact (Eq. *9*). The resulting restoration of a partially-modulated SAM stimulus is shown in Supplementary Fig. 1, while the design of the modified strategy is explained in the Methods (Sec. 7).

The two audio processing types can be applied to any stimulus, with the option of modifying the modulation depth (and dynamic range) of the stimulus envelope or not. Both options aim to improve CS-affected AN processing, but might have qualitative differences in sound-perception outcomes. To generalise the two processing types for different audio stimuli, we first focussed on high-pass (HP) filtered speech (i.e., speech content above *f*_*cut*_ = 1.65 kHz) which relates more to the SAM stimuli. The intelligibility of HP-filtered speech relies on the coding of the temporal envelope of sound [68, 69] and has been linked to the EFR results of listeners in previous CS-related studies [62, 70–72]. After establishing the envelope processing for HP-filtered speech and evaluating the simulated restoration that was achieved, the CS-compensating algorithms can be used in exactly the same way to process any stimulus. Our final goal was to optimally apply our developed strategies to natural, broadband (BB) speech stimuli and improve speech coding in noise, where CS predominantly affects speech perception [9–11, 15, 16].

An example of how our envelope-processing algorithms can be applied to any (modulated) stimulus is shown in Fig. 3a, where the two processing types were applied to a HP-filtered speech-in-noise stimulus using the 13,0,0 CS-profile parameters (Supplementary Table 3). To estimate the modulation envelope and peaks of the noisy stimulus across time, a blind RMS-based procedure was adopted (see Methods). The envelope-processing functions *g* (Eq. *2*) and *g*_*m*_ (Eq.*9*) were computed between the estimated peaks and troughs of the envelope across time, and then multiplied by the noisy stimulus. The processed stimuli (and envelopes) of the two strategies are labelled as 13,0,0 and 13,0,0m, respectively. Fig. 3b shows the restoration yielded by the two algorithms on the simulated AN responses. The original processing strategy (13,0,0) was able to fully restore the CS-affected response peaks back to the NH response by extending the dynamic range, while the modified strategy (13,0,0m) achieved only a partial restoration.

**Fig 3.**
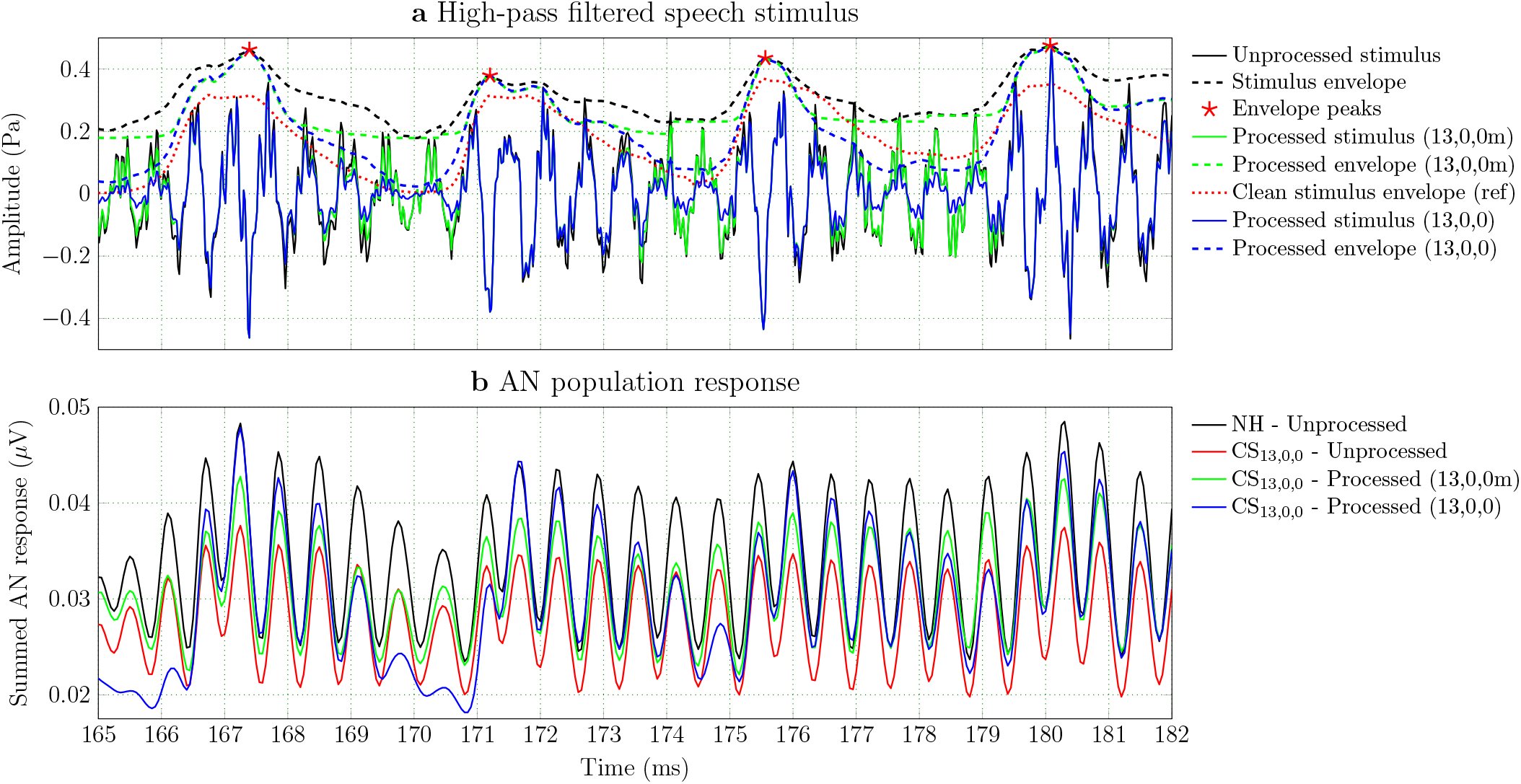
CS compensation of speech in noise. **a** The two processing types were applied for the 13,0,0 CS profile to the vowel /a/ of the word ‘David’, extracted from a sentence of the Flemish Matrix Test [67]. The sentences were high-pass filtered (*f*_*cut*_ = 1.65 kHz) and the generated speech-shaped noise was added to the word with SNR = 0 dB. **b** The unprocessed and processed stimuli were presented 20 times to the model [50], with randomly-selected segments of noise added each time. The unprocessed stimuli were given as input to the NH and CS peripheries, and the processed stimuli to the CS periphery to evaluate the achieved restoration. The simulated AN population responses were then averaged over the 20 stimulus presentations to derive the stimulus-driven responses that are shown here.

Although the algorithms were originally designed for SAM tones, the processing had similar simulated outcomes when applied to HP-filtered speech in noise. In the same way, the algorithms are expected to yield similar restoration when applied to natural, BB speech. Even though our model failed to predict an improvement for natural speech after processing, we still applied our algorithms in our measurements to assess their effect. Finally, to estimate the impact of our CS-compensating algorithms on speech perception, we evaluated their processing using two objective metrics for speech quality and intelligibility, the PESQ [73] and STOI [74], respectively. Supplementary Table 1 shows the average PESQ and STOI scores before and after processing, computed for natural (BB) and HP-filtered Flemish Matrix [67] sentences in noise. To ensure the robustness of our envelope estimates in noise, we also included a reference strategy in which we applied the modified strategies using the estimated envelope of the clean-speech stimulus instead (Fig. 3a; ref). Although the 13,0,0 CS-processing algorithms mostly improved the objective evaluation metrics in BB speech, the 10,0,0 and 7,0,0 algorithms were detrimental to the PESQ and STOI scores, with the original strategies decreasing STOI in both cases. A pilot evaluation of all the processing algorithms in the Flemish Matrix test showed that quality and intelligibility in noise significantly decreased with the 7,0,0 algorithms (results not reported). A more pronounced processing is applied to the 10,0,0 and 7,0,0 compared to the 13,0,0 CS profile (Fig. 2), which results in processed speech sentences that sound less natural and significantly differ from the unprocessed speech. For this reason, and to limit the duration of our tests, we chose to exclude most of the 10,0,0 and 7,0,0 CS-processing strategies from our SI measurements.

## 3 Evaluation

To evaluate the applicability of our algorithms to younger and older listeners with normal audiograms, we assessed possible improvements in physiological as well as sound-perception experiments (Fig. 1b). The evaluation included three measurement sessions (Table 1), in which differences in human responses between processed and unprocessed sound were evaluated using measured EFRs (i.e., EEG potentials evoked by the synchronous response of neurons to the envelope of a stimulus), AM detection sensitivity, and SI. The most relevant types of processing were evaluated in each session based on the assessed auditory test, ensuring that the duration of each measurement was kept under 1.5 hours. The processing algorithms were fast to execute and were applied online to the stimuli of each trial.

**Table 1.** Overview of the measurement sessions. Our measurement protocol was divided into three sessions, each lasting approximately 1.5 hours. For each session, the conditions evaluated are summarised in brackets.

|  |  |
| --- | --- |
| **1st session (EEG)** | SAM tone stimulus (unprocessed, 13,0,0, 10,0,0, 7,0,0)<br>"David" speech stimulus (unprocessed, 10,0,0) |
| **2nd session (AM-SRT)** | AM detection (unprocessed, 13,0,0m, 10,0,0m, 7,0,0m)<br>SRT measurement for BB and HP speech (unprocessed, 13,0,0m-ref) |
| **3rd session (SI scores)** | SI of BB and HP speech (unprocessed, 13,0,0m-ref, 13,0,0, 13,0,0m, 10,0,0m)<br>at 3 fixed SNRs (SRT, SRT-3 dB, SRT+3 dB) |

NH volunteers aged between 18-25 (yNH) and 45-65 (oNH) years old participated in the study and listened to the processed and unprocessed audio for each test. Based on previous reports [11, 14, 75–77], we expected that the older listeners suffered from age-related CS, with correspondingly smaller EFR markers. For all subjects, audiometric thresholds were *≤* 20 dB for frequencies up to 4 kHz, except for four oNH subjects, who are indicated with a darker colour in all the following figures (Supplementary Fig. 2). These four subjects suffer from presumed age-related CS in addition to their OHC deficits.

### 3.1 EEG measurements

During the EEG measurement session, we assessed whether our CS algorithms yielded EFR enhancements in response to SAM tonal stimuli (*f*_*c*_ = 4 kHz, *f*_*m*_ = 120 Hz, *m* = 100%). Four conditions were measured, corresponding to the unprocessed stimulus and the processed versions of the stimulus after applying the processing schemes for the three CS profiles (13,0,0, 10,0,0 and 7,0,0). To evaluate the applicability of our processing strategies to speech, we also assessed whether our algorithms yielded EFR enhancements to a speech stimulus. We used a word from the Flemish Matrix test [67] that was HP filtered and that had a constant fundamental frequency *f*_0_ *≈* 220 Hz. EFRs were recorded for the unprocessed stimulus and a processed version. To limit the duration of the EEG measurement session, only one processing condition was chosen for the speech stimulus (10,0,0). More details about the EEG recording and analysis procedures can be found in the Methods (Sec. 7).

To demonstrate the EFR analysis procedure and create an expectation of what the recorded responses would look like, the EFRs of both stimuli were also simulated using our model [50]. Figures 4a,b show the frequency-domain representations of simulated EFRs to the unprocessed and processed stimuli, as presented to the NH periphery and the corresponding CS-affected periphery. Figures 4c,d also show recorded mean EFRs of one participant (yNH_01_) for the same stimuli. For each condition of the simulated and recorded EFR spectra, the EFR_summed_ magnitude was computed using Eq. *10* to quantify the changes in the EFR magnitude after processing. The recorded EFRs corroborated the simulated enhancement trends, while the simulated responses of Fig. 4a,b suggested that the processed stimuli can also yield EFR enhancements in pristine peripheries (dashed lines). This observation follows our expectations to a certain degree, since we designed our algorithms to optimally drive the (remaining) ANFs. Consequently, our algorithms can enhance peripheral coding in any periphery, possibly resulting in greater benefits for young listeners than for older listeners with less AN fibres.

**Fig 4.**
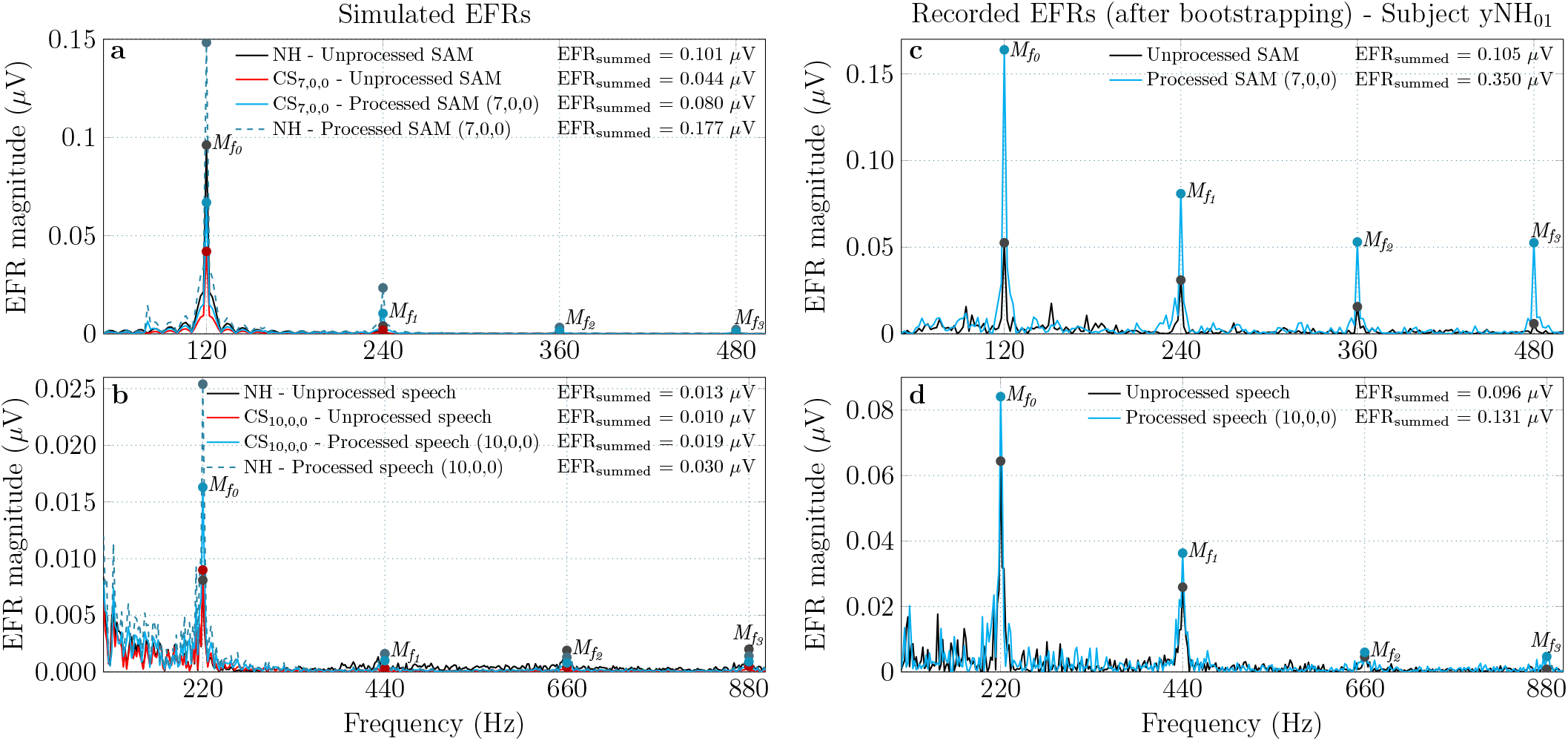
Comparison of simulated and recorded EFRs. **a**-**b** Simulated EFRs to the SAM tone stimulus (**a**) and the HP-filtered speech stimulus (**b**), shown for a NH periphery and a CS periphery in response to both the unprocessed and processed stimuli. **c**-**d** Recorded EFRs of one yNH participant (yNH_01_), in response to the SAM stimulus (**c**) and the speech stimulus (**d**) before and after processing. In all panels, the EFR spectral peaks at the fundamental frequency of the stimulus and the next three harmonics are indicated and were used to compute the EFR_summed_ magnitudes (Eq. 10).

### 3.2 AM detection

During the AM measurement session, we assessed whether the CS algorithms improved behavioural AM-detection sensitivity. The aim was to evaluate whether peripheral coding improvements would also benefit sound perception when adopting the same stimuli. After determining the standard AM detection thresholds (i.e., the minimal modulation depth required to distinguish a modulated tone from a pure tone) using unprocessed stimuli, the stimuli were processed to measure the AM detection thresholds for the three CS profiles. To this end, the modified processing strategies 13,0,0m, 10,0,0m and 7,0,0m were used, to preserve the modulation depth of the AM tones. For each of the three CS conditions, the processing was applied to the modulated tone to yield a stimulus with the same modulation depth but a “sharper” envelope. Because the envelope-processed tones occupied broader FFT spectra after processing, with frequency components beyond the critical-band limits (*±*1 ERB), increased off-frequency listening was expected and may have introduced additional perception cues [78, 79].

### 3.3 Speech intelligibility assessment

During the speech intelligibility (SI) session, we assessed whether the CS algorithms improved SI for natural (BB) and HP-filtered speech. The Flemish Matrix sentence test [67] was used to derive the SRTs (i.e., the SNR required to reach 50% word recognition) of each participant for the two speech types. The SRTs were recalculated after applying one processing algorithm (13,0,0m-ref) to the noisy sentences of each trial, to roughly assess whether the processing improved the SRTs of BB and HP speech. We selected the clean-envelope 13,0,0 modified processing strategy (13,0,0m-ref) for this test, since it gave the best overall objective scores for the processed stimuli (Supplementary Table 1).

Lastly, we individually assessed the benefit to SI based on the measured SRT values of each listener. The aim was to directly compare the effect of each processing strategy on the word-recognition performance of the listeners near their SRT level. It should be noted that the Flemish Matrix test has an intrinsic SRT variability of *±*0.5 dB [67], which could compromise small SI improvements after processing. Therefore, we used fixed SNRs, for which we measured the percentage of correctly recognised words before and after processing (see Methods). We chose 3 SNR conditions corresponding to the SRT, SRT-3 and SRT+3 dB levels of each participant, respectively, measured for both BB and HP speech. As noted above, we limited our experimental setup by only considering processing strategies that were not detrimental to the objective SI evaluation metrics (Supplementary Table 1). To this end, four processing conditions were evaluated here: the two 13,0,0 processing strategies (13,0,0 and 13,0,0m), the modified 10,0,0 strategy (10,0,0m), and the reference clean-envelope 13,0,0 condition (13,0,0m-ref). For each of these four processing conditions and the respective unprocessed condition, the average WRS was measured for every subject for each SNR and speech type.

## 4 Results

The results of our physiological and behavioural measurements (EFRs, AM detection, SRTs and SI percent scores) for the yNH and oNH listener groups are shown in Figs. 5-8. To facilitate a visual comparison between the processed conditions, the median values of the unprocessed conditions for the results of the two groups are indicated by dashed lines in each figure. Blue colours are used for the results of yNH subjects and red for oNH subjects.

**Fig 5.**
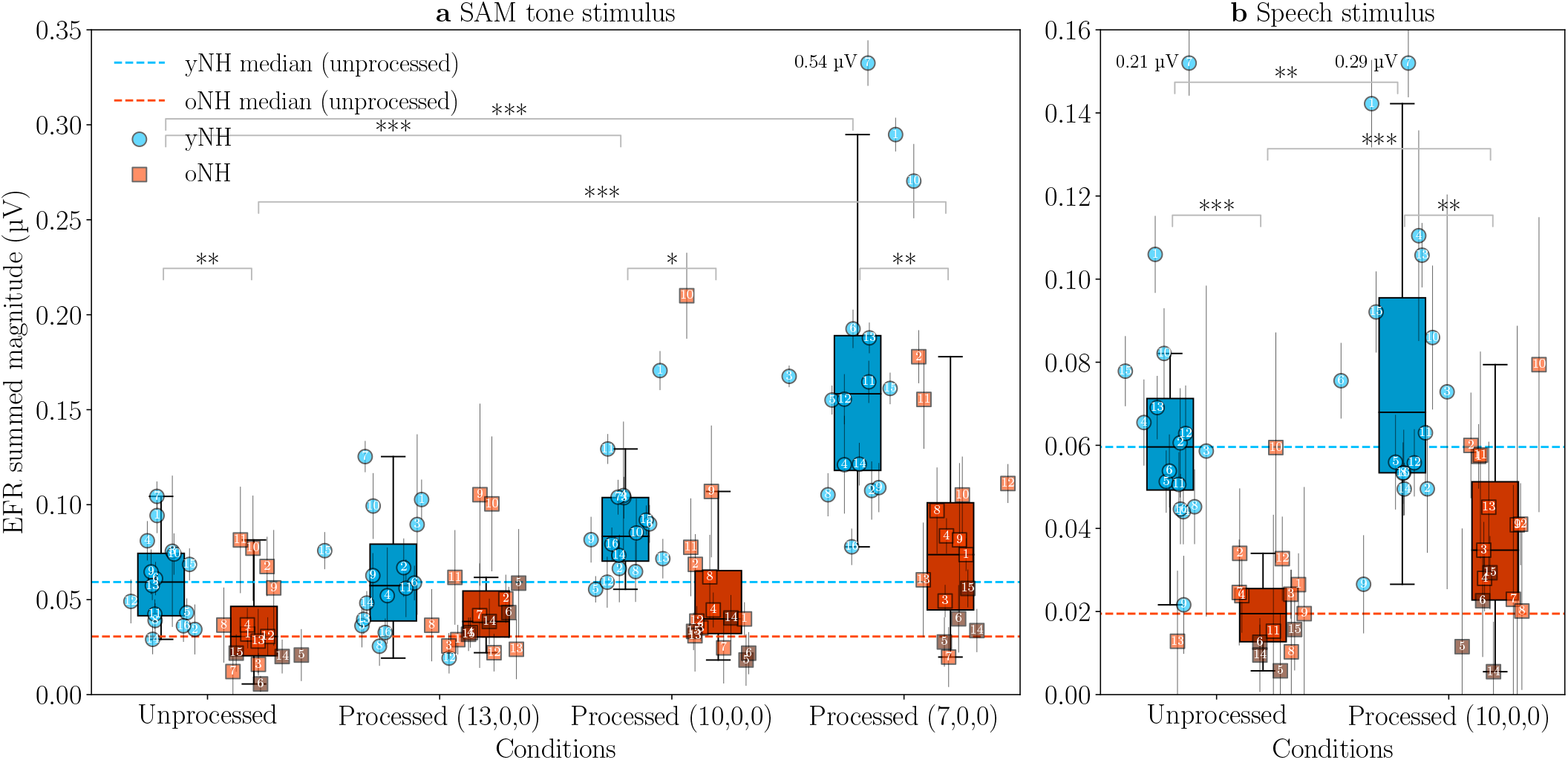
Effect of CS compensation on physiological EFR magnitudes. **a** Recorded EFR results for the SAM stimulus, before and after processing with the three CS algorithms. **b** Recorded EFR results for the speech stimulus (‘David’) before and after processing. In both panels, the individual points correspond to the mean and standard deviation of the summed EFR magnitudes (EFR_summed_), computed from the 400 bootstrapped EFR spectra of each participant (Eq. 10). The box and whisker plots indicate the EFR medians and quartiles of the two groups under each condition

For Figs. 5-7, the effect of age difference between the yNH and oNH groups was assessed using an independent two-sample t-test, computed between the group results for each condition. Statistically significant differences with p-values *<*0.05 are indicated with one asterisk, p-values *<*0.01 with two asterisks, p-values *<*0.001 with three asterisks and p-values *<*0.0001 with four asterisks. Similarly, the effect of processing was assessed separately for each age group using a dependent t-test, by comparing each processed condition to the respective unprocessed result of each group (Figs. 5-8). The t-test results are reported in Supplementary Table 4 and a detailed description of the statistical analysis can be found in Methods (Sec. 7).

### 4.1 EFRs increased after CS-compensating processing

Figure 5 shows individual and median EFR magnitudes (Eq. *10*) for the two subject groups, computed from the EFR spectra of the different conditions. For both unprocessed stimuli, the oNH subjects had lower EFR magnitudes than the yNH subjects (SAM: *p* = 0.0083; speech: *p* = 0.0005), consistent with our expectations of age-related CS for the older group [11, 14, 76, 77]. The 13,0,0 processing condition enhanced the recorded SAM EFR magnitudes for the oNH group (Fig. 5a), but this enhancement was not statistically significant. At the same time, the spread of individual EFR magnitudes within the yNH group was increased by processing, with an interquartile range (IQR) increase from 0.033 µV (unprocessed) to 0.04 µV (13,0,0).

The 10,0,0 and 7,0,0 conditions, however, significantly improved the EFRs in both groups (Fig. 5a), with the 10,0,0 condition showing a statistically significant benefit for the yNH group only (*p* = 0.0001) and the 7,0,0 showing statistically significant differences for both groups (yNH: *p* = 0.0001; oNH: *p <* 0.0001). Statistically significant differences were also found between the performance of the two groups (unprocessed, 10,0,0 and 7,0,0 conditions), demonstrating that there was an effect of age (or age-related synaptopathy) on the magnitude of the recorded EFRs irrespective of the processing applied. The oNH median in the 7,0,0 condition surpassed the yNH median in the unprocessed condition, showing a recovery of the EFR magnitudes in the oNH group to the level of the yNH group after processing. Overall, the SAM EFR processing benefit was more pronounced for the yNH group than for the oNH group, with a median increase of 0.099 µV and 0.043 µV for the yNH and oNH groups in the 7,0,0 processing condition, respectively. At the same time, the 7,0,0 condition significantly spread the individual EFR results (within both groups), with the IQRs increasing by 0.038 µV (116.1%) for the yNH group and by 0.03 µV (116.9%) for the oNH group. This shows that some subjects benefitted more than others from the processing, with those who had unprocessed EFR values below the group median benefitting the least. The estimated EFR noise-floor components did not differ significantly across the measured conditions but showed individual differences among the oNH group (Supplementary Fig. 3), which could explain why some oNH participants did not benefit from the processing..

For the speech-evoked EFR (Fig. 5b), the 10,0,0 condition yielded statistically significant EFR improvements in both groups (yNH: *p* = 0.0073; oNH: *p* = 0.0005). The EFR magnitudes between the two groups also showed statistically significant differences (*p* = 0.0005 for the unprocessed condition). Since the speech-evoked EFR reflects the neural encoding of the pitch information of speech [72, 80], the decreased EFR magnitudes in the older group suggest that pitch tracking was degraded by age-related CS. Although the oNH EFR magnitudes were improved after processing, the (10,0,0) processed speech stimuli were not able to fully restore the oNH magnitudes to the yNH level. Once again, we report larger dispersion (spread) after processing, with the IQRs increasing by 0.02 µV (91.2%) for the yNH group and by 0.016 µV (122.2%) for the oNH group.

### 4.2 Behavioural AM detection thresholds improved after CS processing

Figure 6 shows individual and median AM detection thresholds of the yNH and oNH groups, before and after processing. Both age groups performed similarly for the unprocessed condition (median thresholds of -19.6 dB for the yNH group and -19.2 dB for the oNH group), which corroborates the findings of previous studies that have shown no significant differences between the AM detection performance of subjects with and without suspected CS [81, 82]. All processing schemes improved the median AM thresholds and had a larger effect on the yNH group. The improvements after processing were statistically significant in all cases, except for the oNH group and the 13,0,0 condition, and the benefit increased as the processing compensated more strongly for CS. The 7,0,0 condition significantly improved the AM thresholds, with a 12.5 dB improvement for the yNH group (*p <* 0.0001) and an 8.1 dB improvement for the oNH group (*p <* 0.0001).

**Fig 6.**
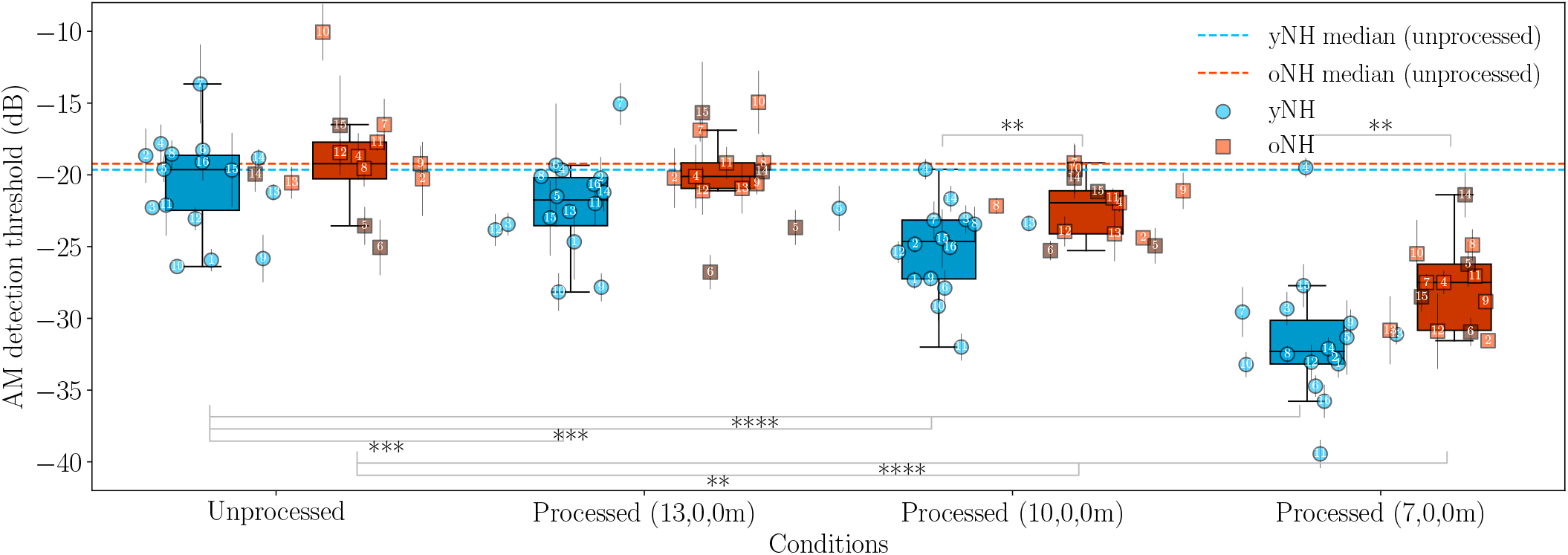
Effect of CS compensation on behavioural AM detection thresholds. For each condition, the mean and standard deviation of the individual AM detection thresholds are shown, computed from three trials for each participant and each condition. The box and whisker plots correspond to the AM-detection-threshold medians and quartiles of the two groups. More negative thresholds indicate better performance.

Statistically significant differences were also found between the performance of the two groups for the 10,0,0 and 7,0,0 processing conditions. It is noteworthy that although both age groups performed similarly for the unprocessed condition, the performance of the groups differed significantly after processing (median thresholds of -32.3 dB for the yNH group and -27.5 dB for the oNH group for the 7,0,0 processing condition). Once again, this shows that yNH subjects benefitted more from the CS-compensating algorithms than oNH subjects.

### 4.3 Mixed outcomes in SI benefit after CS compensation

Figure 7 shows individual and median SRTs for natural (BB) and HP-filtered speech in the oNH and yNH subject groups. In contrast to the observed EFR and AM detection processing benefits (Figs. 5,6), the SRTs were degraded after processing, especially for the oNH group. A small but insignificant SRT improvement was observed for the BB condition, but only among the yNH group, with a median decrease of 0.2 dB from the unprocessed BB SRT. For HP-filtered speech (which relies on temporal-envelope processing), the yNH SRTs had an almost identical median but showed a larger dispersion after processing, with an IQR increase from 1.3 dB to 3.3 dB. A statistically significant difference was also observed between the performance of the two age groups in HP-filtered speech (*p* = 0.0274), but not in BB speech. This corroborates our hypothesis suggesting degraded temporal-envelope coding in the older group, since only the high-frequency component of speech was less intelligible for these subjects. Although consistent improvements were not found for SRTs after processing, Fig. 7 shows that there were several yNH subjects who benefitted from processed speech under both speech conditions (e.g., yNH_05_, yNH_14_, yNH_15_). These subjects corresponded to data points that were below the group median, i.e., subjects who had the best speech intellibiligity performance (lowest SRTs) in the unprocessed condition.

**Fig 7.**
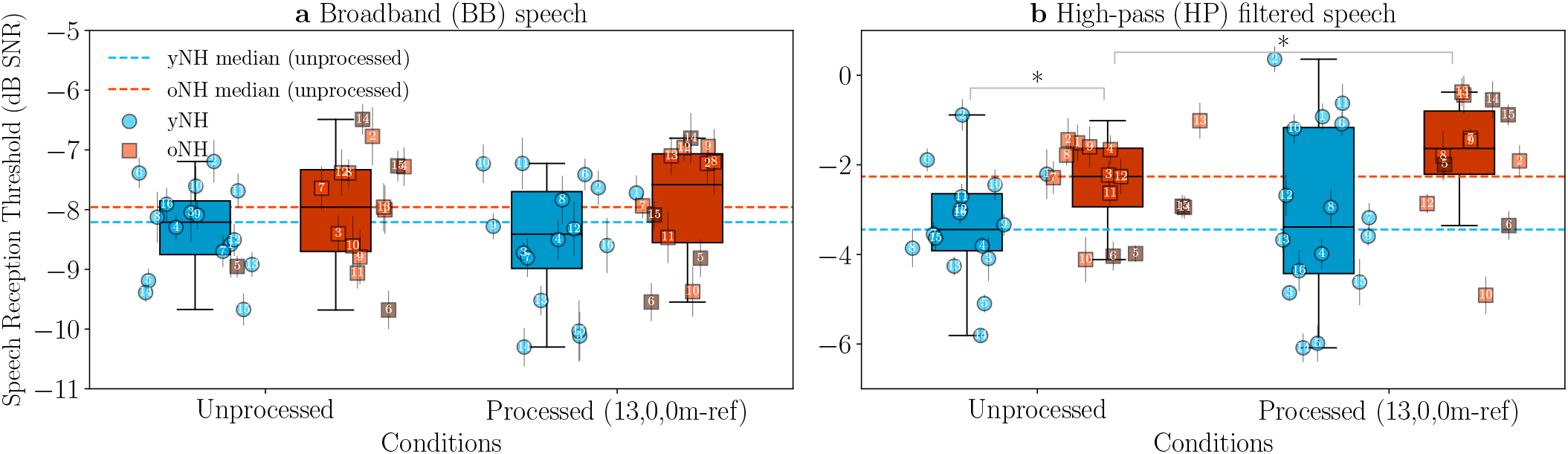
Effect of CS compensation on behavioural speech-reception thresholds. **a** Measured SRTs for BB speech before and after processing with the clean-envelope 13,0,0 modified processing strategy (13,0,0m-ref). **b** Measured SRTs for HP-filtered speech before and after processing with the 13,0,0m-ref processing strategy. For each participant and each condition, the mean and standard deviation was computed from the last 6 reversals of the respective SRT measurement. The box and whisker plots correspond to the SRT medians and quartiles of the two groups. More negative SRTs indicate better performance

Finally, Fig. 8 shows the SI scores (WRSs) before and after processing, computed for BB and HP-filtered speech at three SNRs (SRT, SRT-3, SRT+3 dB). Since the SNR levels (at which the scores were computed) were individually adjusted to the SRTs of each participant for BB and HP-filtered speech (Fig. 7), both group medians were expected to lie at *∼*50% for the BB (SRT) and HP (SRT) unprocessed conditions (Fig. 8b,e). The SNRs of *±*3 dB SRTs (Fig. 8a,d and Fig. 8c,f) corresponded to two different points on the psychometric functions of the participants, and resulted in measured median scores of *∼*14% and *∼*32% for BB and HP-filtered speech at SRT-3 dB, respectively, and to *∼*83% and *∼*73% for BB and HP-filtered speech at SRT+3 dB, respectively. Assuming that the *±*3 dB deviation from the individual SRT resulted in points on the linear part of the psychometric function [67], the average slope of the curve was 11.3 *±* 1.5 %/dB for BB speech and 6.5 *±* 0.1 %/dB for the HP-filtered speech.

**Fig 8.**
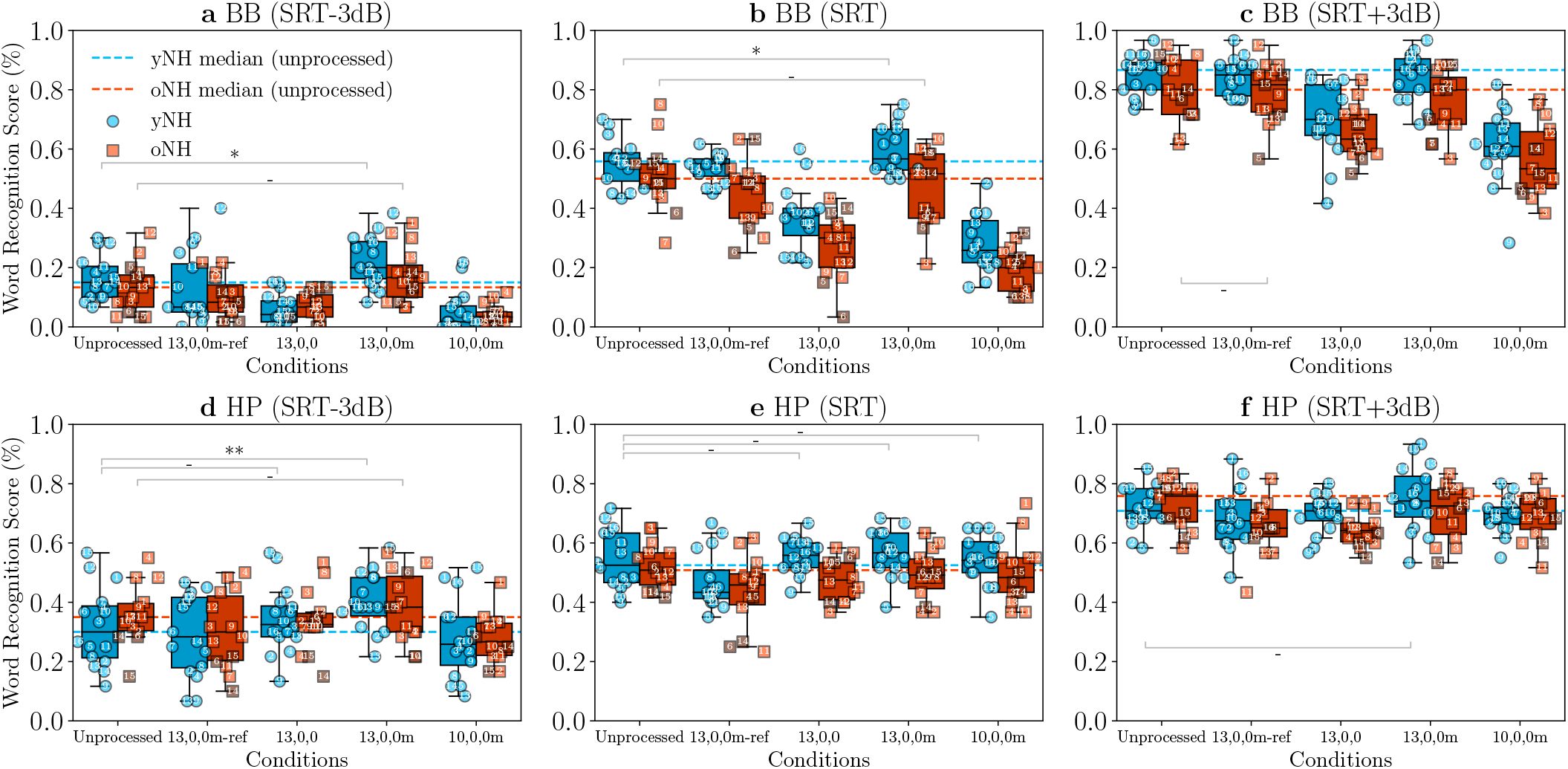
Effect of CS compensation on word-recognition scores near the individual SRT. For BB and HP-filtered speech at 3 SNR levels (SRT-3, SRT, SRT+3 dB), the average WRS of each participant was computed for each condition from 12 measured trials. Statistical significance between unprocessed and processed results is shown only for the cases where a higher score median was obtained after processing. The box and whisker plots correspond to the score medians and quartiles of the two groups. A higher score indicates better intelligibility

For natural (BB) speech (Fig. 8a-c), the 13,0,0 and 10,0,0m processing conditions mostly decreased the individual SI scores of participants, with a less pronounced effect for the HP-filtered speech condition (Fig. 8d-f). For the 13,0,0 modified conditions (13,0,0m-ref and 13,0,0m), a similar effect as in Fig. 7 was observed, with the processing mainly improving the performance of yNH participants. The 13,0,0m processing resulted in improved median scores for yNH participants among all SNR conditions, and to small improvements for oNH participants in some cases. For the most difficult SNR condition (SRT-3 dB), the median SI increased by 5% for the yNH group (*p* = 0.0155) and by 3.3% for the oNH group (*p* = 0.0824) in BB speech (Fig. 8a), and by 8.3% for the yNH group (*p* = 0.0028) and 3.3% for the oNH group (*p* = 0.5121) in HP-filtered speech (Fig. 8d). Based on the estimated psychometric curve slopes of the BB and HP conditions, this corresponds to an SRT improvement of 0.4 dB and 0.3 dB for the yNH and oNH group, respectively (natural speech), and to an SRT improvement of 1.3 dB and 0.5 dB for the yNH and oNH group, respectively (HP-filtered speech).

In contrast to the 13,0,0m processing condition, the clean-envelope reference condition (13,0,0m-ref) did not show consistent improvements to the SI scores. Thus, processing based on a blind approximation of the noisy envelope was more robust than providing the envelope of the clean stimulus. The median differences between the 13,0,0m and 13,0,0m-ref processing conditions were more pronounced for lower SNRs (Fig. 8), and this suggests that enhancing the modulations of the stimulus the participants listened to (noisy speech) was more important for this type of processing.

### 4.4 Who benefits from CS-compensating processing?

To investigate the characteristics of those who benefitted more strongly from CS processing, we studied the relationships between the different physiological and behavioural measures using Spearman’s rank-order correlation *r*. First, correlations were computed between measured EFR magnitudes, AM detection thresholds, SRTs, and audiometric thresholds. Supplementary Tables 5-8 show the values of Spearman’s *r* between the SAM EFR, speech EFR, AM detection and SRT results for the cases where at least one statistically significant correlation was found. For each metric, correlations were also computed for the improvement of each processed condition, i.e., after subtracting the respective unprocessed measures from each processed condition. Two metrics were considered for the correlations relating to audiometric thresholds: the hearing threshold at 4 kHz and the average extended-high-frequency (EHF) hearing thresholds (10-16 kHz). Finally, Supplementary Table 9 summarises the statistically significant correlations of the WRS improvement for all processing algorithms, computed as the SI improvement of each processing condition compared to the unprocessed SI score. Age was not included as a variable in the statistical analysis due to the large age differences between the two groups, and we found no statistically significant correlations between age and the remaining metrics when examining each group separately.

#### Correlations between EFR magnitudes and behavioural AM detection thresholds

Strong and statistically significant correlations were found between AM detection thresholds and EFR magnitudes (Supplementary Table 7), especially for the 7,0,0 processing. Figure 9c shows the most pronounced relationship among the measures, computed between 7,0,0m-processed AM detection thresholds and 7,0,0-processed SAM EFR improvement from the unprocessed condition (7,0,0 - unprocessed). All correlations were negative (*r <* 0), showing that AM thresholds generally decreased as the EFR magnitudes, or as the improvement of the EFR magnitudes after processing, increased (and vice versa). This indicates a strong link between physiological and behavioural metrics when the same stimuli and processing methods are adopted. Although similar relationships were found for each age group separately (Fig.9a,b), the correlations were not statistically significant in these cases.

**Fig 9.**
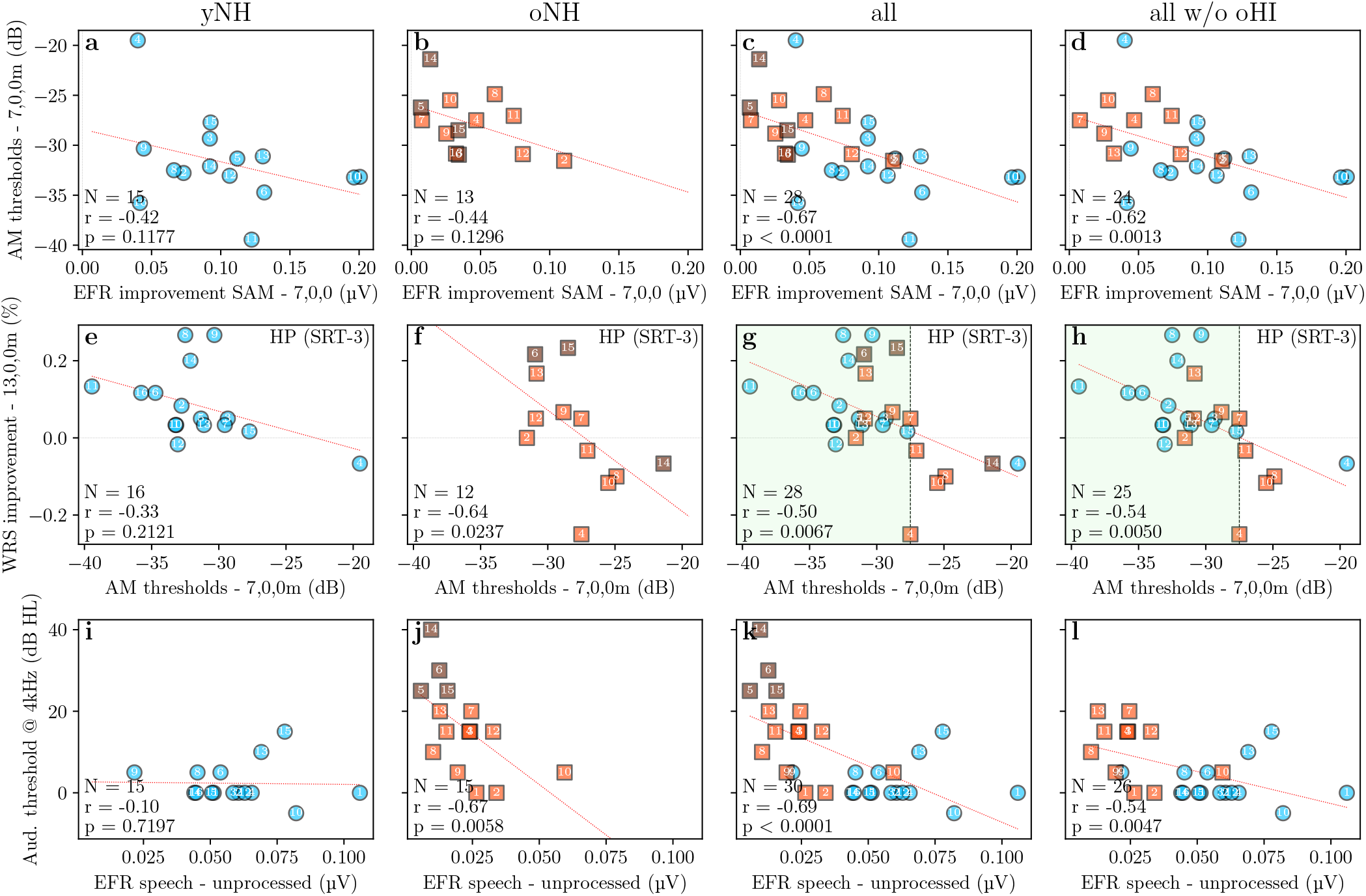
Significant correlations between metrics. **a**-**d** Correlations between the SAM EFR improvement after processing with the 7,0,0 condition (7,0,0 - unprocessed) and the AM detection thresholds of the 7,0,0m condition. **e**-**h** Correlations between the AM detection thresholds of the 7,0,0m condition and the WRS improvement after processing with the 13,0,0m condition (13,0,0m - unprocessed), computed for HP-filtered speech at SNR = SRT - 3 dB. **i**-**l** Correlations between the EFRs to the (unprocessed) speech stimulus and the measured audiometric thresholds at 4 kHz. Correlations are shown between each group separately (yNH and oNH), for all subjects together (all), and for all subjects after removing the oNH subjects with audiometric thresholds *>*20 dB HL (all w/o oHI). In each panel, the sample size *N* and the values of the Spearman’s rank-order correlation *r* and the probability *p* are shown on the lower left corner. The red dotted lines correspond to the linear regression model fit to the data.

#### Correlations with SI

Overall, the most significant correlations to the measured SI were found for HP-filtered speech (Supplementary Tables 7,9). The metric that showed most correlations to the measured SI of the participants was the AM detection performance (Supplementary Tables 8,9). For the lowest SNR (SRT-3 dB), AM detection thresholds in the 7,0,0m condition showed negative correlations to the WRS improvement after processing (Supplementary Table 9). Thus, participants with good AM detection performance (low thresholds) benefitted more from the processed speech than those with poor AM sensitivity. As an example, Fig. 9g shows the relation between the 7,0,0m AM thresholds and the WRS improvement of HP-filtered speech at the noisiest scenario. Most participants with AM thresholds below -27.5 dB (light green part) were able to benefit from the 13,0,0m processed speech. Hence, the AM detection performance in the 7,0,0m condition could provide an indication of the expected SI score improvement after CS processing. The correlation was more evident in the oNH group (Fig. 9f), and became stronger when the oNH subjects with elevated thresholds were not considered (all w/o oHI).

#### Correlations with audiometric thresholds

When examining both age groups together, EFRs and AM detection thresholds correlated well with the audiometric thresholds at 4 kHz and at the EHFs. Statistically significant negative correlations were found between audiometric thresholds and measured EFRs (Supplementary Tables 5,6), especially for the 7,0,0-processed SAM EFR condition and the speech EFR (unprocessed). Thus, participants with more elevated audiometric thresholds (at 4 kHz and EHFs) had lower EFR magnitudes, as illustrated in Fig. 9k for the 4 kHz audiometric thresholds. At the same time, statistically significant positive correlations were also found between audiometric thresholds and measured AM detection thresholds for the 7,0,0 condition (Supplementary Table 7). This suggests that people with OHC deficits performed worse on this task after processing, in line with the results of Fig. 6.

However, when removing the oNH subjects with elevated thresholds from the statistical analyses (all w/o oHI), the statistical significance of the audiometric-threshold correlations consistently decreased (Supplementary Tables 5-9). This is more visible in Fig. 9l, where the *p* value significantly increased after removing the subjects with elevated thresholds, indicating that these four subjects could have biased the statistical analysis (Fig. 9j). At the same time, almost no statistically significant correlations to the audiometric thresholds were found when examining the yNH and oNH groups separately (Supplementary Tables 5-9). This suggests that the age difference between the two groups is the most important factor here and not the audiometric thresholds per se, which also implies the effect of age-related CS [6, 59, 75, 76, 83]. Nevertheless, subjects with HL at high frequencies were expected to have decreased responses to HP-filtered stimuli (EFR speech) and to perform worse at the detection of high-frequency tones (AM task).

## 5 Discussion

Our data show that physiological EFRs and perceptual AM sensitivity were enhanced in both yNH and oNH listeners when using our CS-compensating algorithms. The “mildest” CS-processing condition (13,0,0), which focussed on MSR and LSR fibre loss only, did not yield consistent improvement for all subjects, but the 10,0,0 and 7,0,0 conditions systematically improved EFR magnitudes and AM thresholds. The latter two conditions were designed to compensate for selective loss of all ANF types and applied more pronounced processing to the stimulus envelope. Based on the outcomes of several animal studies that have shown a preferential loss of LSR and MSR ANFs with CS (e.g., [3, 5, 84]), we selected to eliminate these fibre types in our CS-compensating algorithm design. At the same time, it has been argued that LSR and MSR fibres are not crucial for the coding of sound at moderate-to-high levels [85], which further justifies our focus on compensating for losses of HSR ANFs instead. However, a complete loss of LSR and MSR fibres with CS is not certain [66], considering also that synaptic regeneration is possible after noise exposure (as has been reported in mice and guinea pigs [84, 86, 87]). Thus, more uniform losses of the different ANF types can be used in the algorithm design of future CS-compensating strategies and be evaluated experimentally.

Furthermore, statistically significant correlations were found between AM detection thresholds and SAM EFR magnitudes (Supplementary Table 7, Fig. 9c), showing that the AM detection performance of a subject (especially in the 7,0,0m condition) can predict the strength of the recorded EFR to the same stimulus paradigm (and vice versa). In line with the rectangular-wave-amplitude-modulated (RAM) stimulus paradigm [14], our envelope processing spread out the frequency components of the AM tonal stimuli we used in this study. This could have caused additional perception cues when listening to the processed sounds [78,79] and could partially explain the improvements of both EFR and AM-detection metrics [14]. Speech recognition in the Matrix sentence test showed a small, but inconsistent, improvement across participants after processing: The yNH group and those with high AM detection sensitivity benefitted the most from processed speech (Fig. 9g). The latter aspect goes in line with the findings in [19, 45], where individuals with AN deficits and poor modulation detection sensitivity performed worse in speech recognition. At the same time, since our hearing-enhancement algorithms were developed to compensate for the functional effects of CS, they yielded processing that optimally drives the ANFs. Thus, even though the algorithms were designed to optimally restore CS-affected responses to NH, the young listeners could have benefitted more from processing due to the increased number of fibres compared to the older listeners (e.g., Fig. 4). Additionally, older listeners could have had mixed pathologies (together with CS) which were not accounted for in this work.

The evaluated processing strategies had individual effects on the SI benefit, with the 13,0,0 modified strategy (13,0,0m) yielding overall the best WRS results (median improvement of up to 5% and 8.3% for the yNH group in the recognition of BB and HP-filtered speech, respectively). Although the 13,0,0 modified strategy improved intelligibility in most cases (Fig. 8), similar benefits were not encountered for the same processing type when applied for the 10,0,0 profile (10,0,0m). This goes in line with the objective evaluation and our pilot assessment (Sec. 2.4), where the 10,0,0 and 7,0,0 strategies further decreased intelligibility for speech in noise. At the same time, the 13,0,0m strategy outperformed the 13,0,0 strategy, even though the simulated AN response enhancement was lower in the first case (e.g., Fig 3). Thus, perceptual markers are still necessary in the algorithm design (e.g., in terms of speech intelligibility and quality) to objectively evaluate the CS-compensating strategies alongside the simulated AN response restoration.

To this end, we chose to use the PESQ and STOI metrics for each processing strategy (Supplementary Table 1). The STOI evaluation resulted in higher scores for the modified strategies in comparison to the original strategies, which was in line with our measured SI results (when comparing the 13,0,0m to the 13,0,0 SI scores). Although our simulations suggested that the modified strategies can achieve lower CS restoration, this type of processing preserved the envelope modulation depth and resulted in smoother and more “natural-sounding” speech stimuli than the original processing strategies, which could be the reason for the higher STOI scores. On the other hand, the highest PESQ scores were achieved by the original processing strategies (BB speech), which could be justified by the possible noise suppression that these might achieve as a side effect. Supplementary Table 1 also shows that the clean-envelope 13,0,0 modified strategy (13,0,0m-ref) gave the best scores by improving both PESQ and STOI in BB speech after processing. However, this strategy showed the worst SI results for our subjects, without showing consistent improvements to the SRTs and the WRSs or significant correlations to other metrics. Thus, the evaluation of augmented speech using such objective metrics might be unsuitable, and this may relate to the fact that the PESQ and STOI metrics consider the clean-speech signal as the (ideal) reference. Processed speech would ideally require an evaluation approach that can connect the simulated AN response restoration to predicted intelligibility for the individual HI periphery. To this end, the integration of a backend in the auditory periphery model that can simulate (psychoacoustic) speech-recognition performance [88], or the use of similar human-inspired frameworks [89] might prove helpful in future studies.

Although we performed a short training at the beginning of each SI session (see Methods), stronger (or more consistent) SI improvements may be observed when allowing the subjects to get accustomed to the processed sound for a more extended time period [41]. The results of the last measurement session (WRS) were expected to reflect the performance of listeners that were already more familiar with the processed sounds (taking into account the previous SRT session), but a longer training session or a shorter interval between the sessions might still be necessary. This is better demonstrated in Supplementary Table 10, where the median WRS improvement was computed for our best-performing strategy (13,0,0m) in the two groups. Overall, the SI improvement was more significant after omitting the first four trials of the results (up to 10 % improvement in the noisiest scenario), hence there still seems to be a training effect over the first trials of the session that should be accounted for in future studies.

The observed discrepancy between the improvements in peripheral coding (EFRs, AM detection) and in SI after processing might be attributed to the existence of compensatory mechanisms in the auditory pathway after the AN, e.g., in the auditory brainstem [90], midbrain [91] or cortex [92]. This may point to a general limitation of current auditory signal-processing treatments for ANF damage, thus future SNHL compensation strategies might benefit from accounting for central auditory-processing changes that follow CS (e.g., central gain), with respect to the effect of such compensatory mechanisms on speech coding as such. At the same time, the distortion of speech after processing could also be a reason for the individual differences and small benefits in SI scores. Although our algorithms were optimised purely on the basis of the simulated outcomes of a biophysical model [50], additional signal-processing techniques could be applied after the optimisation procedure to minimise the negative effects that envelope processing might introduce (e.g., [37, 39, 44, 47, 93, 94]). The presented algorithms rely on instantaneous processing of the temporal envelope (RMS window of *<*1 ms), which has been argued to lead to non-linear distortion and poor sound quality in dynamic-range compression [17, 32, 34, 35, 37, 94–97]. Thus, when processing speech, a release time constant can be added to the processing algorithms to reduce distortion [97, 98], in line with standard slow-acting compression algorithms. It should also be noted that our algorithms further emphasise temporal amplitude fluctuations by increasing the intensity contrasts and modulation depth of speech (without applying any gain), and could thus limit the adverse effects of noise on speech perception [18, 97, 99]. This is in constrast to standard fast-acting compression that is known to reduce the amplitude fluctuations of brief speech components and can distort temporal cues [17, 18, 97, 100]. On the other hand, our processing was applied on the envelope of the broadband signal, assuming CS pathologies that are equally distributed across frequency. Previous studies have reported that processing the whole speech envelope can lead to perceivable speech distortion [35, 37]. Thus, a possible extension of the proposed algorithms would be to process the envelope in more frequency bands (multi-channel processing [17]), to target the enhancement of both spectral and temporal contrast of the envelope modulation [99, 101], and also to compensate for frequency-specific types of CS and/or OHC loss, depending on the individual SNHL type.

The design of our CS-compensating processing function focussed most strongly on stimulus envelope regions that fluctuate the most, i.e., where temporal modulations (intensity contrasts) of the signal are the strongest. As a result, this type of processing mostly improves the voiced parts of speech stimuli, resulting in more enhanced vowels and voiced consonants than unvoiced consonants. At the same time, modification of the temporal modulation characteristics of speech may attenuate short-duration parts that are low in intensity compared to the rest of the stimulus, such as unvoiced stop consonants or rapid formant transitions between voiced vowels and consonants [39]. This could deteriorate the perception and intelligibility of consonants and introduce difficulties in word identification [37, 102], especially for CS-affected listeners that might additionally suffer from impaired detection of short-duration signals [16]. In future work, the processing schemes can be improved to enhance the low-intensity, short-duration consonant cues more strongly, instead of enhancing long-duration cues such as vowel formants [39], e.g., by increasing the consonant-vowel intensity ratio [38, 46, 99, 103]. A more thorough analysis of our SI results or a more specific measurement session (e.g., monosyllabic word tests) could provide insight regarding the individual intelligibility enhancement of vowels and consonants after processing.

Overall, the processing conditions showed larger dispersion of the individual results for each of the tests (EFRs, AM detection thresholds, SRTs and WRSs). This indicates that the processing has a strong individual component, with the variability increasing as the processing became stronger (from 13,0,0 to 7,0,0). Because the most pronounced processing condition (7,0,0) produced SAM stimuli that are similar to the RAM stimulus [14], the evoked responses are expected to be more sensitive to CS than to OHC deficits [14, 62]. Thus, the individual degree of CS could be a possible explanation for the spread of the physiological and behavioural results as the processing became more pronounced. Listeners with higher degrees of CS (and hence smaller unprocessed EFR markers) may be those who benefitted less from CS-compensating processing, however this was not clear from the results of our statistical analysis. To this end, diagnostic assessments of the individual degree of CS in each participant [58, 59] could help to explain the individual performance differences between participants with more certainty.

## 6 Conclusion

Our proposed hearing-enhancement strategies go beyond conventional hearing-aid algorithms that typically apply dynamic-range compression to compensate for OHC loss. Instead, our type of processing either preserved or extended the dynamic range of the temporal speech envelope, by increasing the dynamic range of the simulated CS-affected responses to compensate for selective losses of ANFs (CS). The envelope processing was designed on the basis of SAM tones and afterwards extended to HP-filtered and natural (BB) speech, targeting the enhancement of temporal modulation in the periodicity envelope. The proposed CS compensation was able to significantly improve peripheral coding in both young and older NH groups, enhancing their EFRs and AM detection performance. The strategies were not able to improve the SI of all participants, but benefitted those with good behavioural performance, especially among the yNH group. For SNRs at 3 dB below the individual SRTs, our best-performing algorithm improved word recognition of speech by 5% in the yNH group (*p <* 0.05) and by 3.3% in the oNH group (n.s.).

Although our proposed hearing-enhancement strategies were based on the compensation of AN responses in CS-affected peripheries, the resulting algorithms were able to improve temporal-envelope processing for listeners both with and without suspected age-related CS. However, when applying our algorithms to speech in noise, intelligibility showed statistically significant improvements for young normal-hearing subjects but not for subjects with suspected age-related CS. Although speech processing might require adaptations to restore hearing in listeners with CS pathologies, augmented hearing could be achieved for NH listeners, allowing for speech-enhancement applications in acoustic scenarios where intelligibility is known to suffer (e.g., train-station announcements, smart phones, hearables). At the same time, the processing algorithms were easy to implement and fast to execute, and were used online in our measurements without adding significant delays between trials. Thus, our proposed type of sound processing may extend the application range of present-day hearing aids and hearables by improving temporal-envelope processing, while leaving sound amplification unaffected.

## 7 Methods

Using a biophysically inspired model of the auditory periphery [50], we applied a numerical method that iteratively altered a sound-processing function so as to minimise the difference between simulated auditory CS responses and NH responses (Fig. 1). In each iteration, the AN population response to the modified stimulus was computed and compared to the reference response, until an error measure was sufficiently minimised. An iterative formula was used:

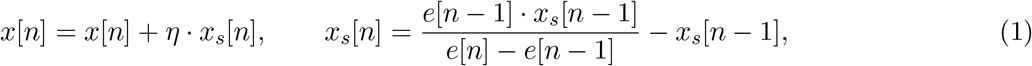

where *n* is the iteration number, *x* the unknown parameter and *x*_*s*_ its respective step, *η* the learning rate and *e* the error measure to be minimised. A learning rate *η* = 0.8 and an initial step *x*_*s*_ = 0.1 were used, and the procedure was repeated for each unknown parameter until the step *x*_*s*_ was lower than 0.01.

### 7.1 Sound processing for fully-modulated SAM tones

The optimisation was performed for 50-ms-long SAM tonal stimuli (carrier frequency *f*_*c*_ = 4 kHz, modulation frequency *f*_*m*_ = 120 Hz and *m* = 100% modulation) of levels from 0 to 120 dB SPL and a step size of 1 dB SPL. Each time, the error measure *e* was defined as the mean-squared error (MSE) between the peak AN population responses, computed at 37-42 ms after the stimulus onset to ensure a steady-state response. The maximum values of the two responses were computed over this region and 10 samples before and after their respective maxima (*±*0.5 ms) were considered for the MSE difference, accounting for possible phase differences between the two responses.

The iterative formula of Eq. *1* was used to independently derive the parameters *a, b* and *c* of a non-linear envelope-processing function *g* for each level

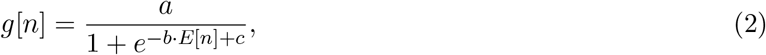

where *E* is the envelope of the input auditory signal *x* and *n* is each time sample. During the optimisation, the Hilbert envelope was computed for the SAM tones, corresponding to the amplitude fluctuations of the modulating tone signal (*f*_*m*_ = 120 Hz). The parameters *a, b* and *c* were initialised at 1, 87.5, and 6.25, respectively. The optimisation procedure was applied twice to ensure that an optimal combination of the three independently-defined parameters was derived. The values of the variables *a, b* and *c* define the shape of the non-linear function: *a* is the variable determining the maximum gain applied, *b* defines the slope ofthe function and *c* the offset (lower operating point) of the function. Although the slope of the non-linear function (variable *b*) was kept constant for each CS profile, different values of the parameters *a* and *c* were computed for each stimulus level to achieve level-dependent processing.

After defining the optimal parameter values for each stimulus level, two exponential functions were fitted to the acquired values of the parameters *a* and *c*. These functions were selected based on the exponential shapes of the derived parameter values across level. The formulae for computing all the parameters of Eq. *2* are given below, expressed as functions of the stimulus-envelope peak. The fitted parameter values used in the formulae are given in Supplementary Table 2.

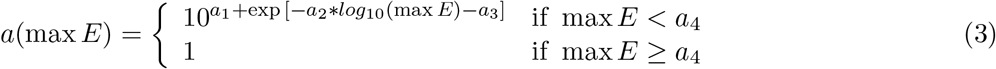

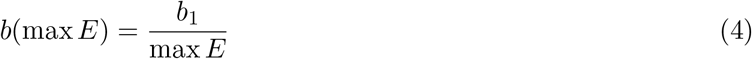

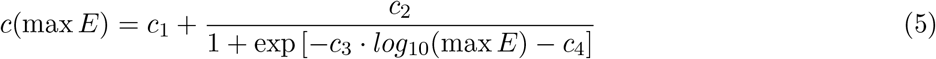

The three formulae can be used to apply level-dependent processing to a SAM tone based on the peak (maximum value) of its envelope, but they can also be adjusted to use the respective RMS value instead. Here, we focussed on applying our processing to 70 dB-SPL stimuli and ensuring that the responses to supra-threshold stimuli can be enhanced after processing. Supplementary Table 3 shows the values of the three parameters for each chosen CS profile at 70 dB SPL. Depending on the CS profile, the respective values of the variables *a, b* and *c* can be used directly in Eq. *2* to compute the non-linear function *g*. The function *g* can then be multiplied by the stimulus to obtain a processed signal that can compensate for the specific CS profile at 70 dB SPL. The parameter values of Supplementary Table 3 were used in the non-linear processing algorithms of each CS profile for our simulated (Sec. 2) and experimental (Sec. 3) evaluation.

### 7.2 Sound processing for partially-modulated stimuli

To apply the processing functions to SAM tones of *<*100% modulation without affecting the modulation depth, we modified the range of the non-linear function to have a different lower operating point. First, the envelope *E* of the modulated signal was scaled to remove the envelope offset (modulation depth):

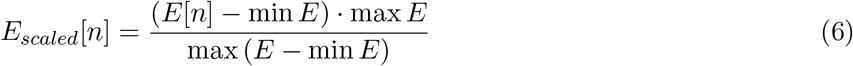

Then, the processing function of Eq. *2* was applied to the scaled envelope *E*_*scaled*_ instead:

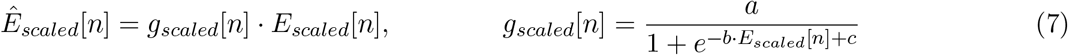

The operation of Eq. *6* was reversed to rescale the processed envelope 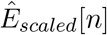 back to the range of the envelope *E* of the original signal:

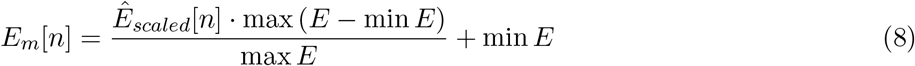

The resulting envelope *E*_*m*_ corresponds to the desired processed envelope, which differs from the original envelope *E* only in the regions between the envelope peaks and troughs. By dividing the derived envelope *E*_*m*_ by the original envelope *E* we get the desired processing function:

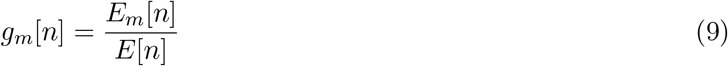

The modified processing function *g*_*m*_ can thus be applied to a modulated signal and preserve the original modulation depth of its envelope (Supplementary Fig. 1). By using the modified function *g*_*m*_, the processing is applied solely to the slope of the modulation envelope. However, as shown in Supplementary Fig. 1, the processed stimuli are unable to fully restore the CS-affected AN responses to the NH responses. As the modulation depth decreases, the processing of the envelope gradually becomes less effective, until the envelope becomes flat (*m* = 0) and our algorithms apply no processing to the signals. It should be noted that in the case of fully-modulated stimuli, both the original and the modified functions apply the same processing.

### 7.3 Speech processing

Evaluation of the algorithms for speech was performed using the material of the Flemish Matrix test [67]. The Flemish Matrix includes 260 sentences recorded by a female speaker, each comprised of a 5-word combination from a closed set of 50 Flemish words. To generate the HP speech material, the (BB) Flemish Matrix sentences were HP filtered (zero-phase digital filtering) using an FIR filter with an order of 1024 and a cutoff frequency *f*_*cut*_ = 1650 Hz (as in [70, 71]). Then, speech-shaped noise (SSN) material was generated from the long-term average speech spectrum of all sentences (as in [67]), separately for the BB and HP-filtered material. In each case, the BB SSN generated was used for the evaluation of BB speech in noise and the HP SSN for the evaluation of HP-filtered speech in noise.

To apply our envelope processing to speech, we performed an automatic RMS-based envelope estimation from the speech waveform using a running RMS window of 25 samples (*∼*0.57 ms for a sampling frequency f_s_ = 44,100 Hz). The estimated envelope was then smoothed using a moving-window function of the same size (25 samples), to remove the high-frequency fluctuations of the envelope and capture the speech pitch periodicity [24]. To find the operating points of the non-linear functions, we used the peaks of the envelope that had an amplitude larger than 1.15 times the amplitude of their neighbouring samples. The corresponding troughs between each pair of peaks were then computed from the local minima of the envelope between peaks, and the envelope was normalised to the local maxima of the signal. Finally, based on the estimated peak and trough positions, the attenuation function *g* or *g*_*m*_ (Eqs. *2,9*) was computed between each pair of troughs using the normalised envelope, and was multiplied by the original stimulus to generate the envelope-processed speech.

The aforementioned process was applied in the same way for quiet or noisy stimuli and was visualised in Fig. 3a for a speech-in-noise stimulus. In the case of stimuli in noise, the two processing functions were directly applied to the estimated envelope of the noisy stimulus without having access to the clean stimulus or envelope (blind processing). However, we also included a reference condition that used the estimated envelope of the clean stimulus (ref). In this case, the envelope-processing function was computed across time from the peaks and troughs of the clean envelope and then applied to the noisy signal to derive the processed version for the desired CS profile (abbreviated as 13,0,0m-ref, 10,0,0m-ref and 7,0,0m-ref for the modified strategy).

### 7.4 Evaluation

For the evaluation of our algorithms, participants were recruited into two groups: 16 young normal-hearing (yNH: 22.5 *±* 2.3 years, 9 female) and 15 older normal-hearing (oNH: 51.3 *±* 4.4 years, 12 female). Volunteers with a history of hearing pathology or ear surgery were excluded as per a recruitment questionnaire. Audiograms were measured in a double-walled, sound-attenuating booth, using an Interacoustics Equinox audiometer. The hearing thresholds of the participants were assessed at 12 standard audiometric frequencies between 0.125 and 16 kHz. The stimuli were presented monaurally to the best ear, determined on the basis of their audiogram and tympanogram. Audiometric thresholds were below 20 dB hearing-loss (HL) for frequencies up to 4 kHz in the yNH group (Supplementary Fig. 2). In the oNH group, audiometric thresholdsn were *≤* 20 dB HL for frequencies up to 4 kHz for all subjects except for four subjects whose results were tagged by a darker colour.

The experimental data were collected in a sound-proof and electrically-shielded booth and the sounds were presented monaurally via Etymotic ER-2 insert earphones connected to a Fireface UCX external sound card (RME) and a TDT-HB7 headphone driver. The EEG recordings were conducted while subjects were sitting in a comfortable chair with a head rest and were watching muted movies. The processing algorithms were implemented in MATLAB and the participants’ psychoacoustic responses were collected using the AFC toolbox for MATLAB [104]. Participants were informed about the experimental procedure according to the ethical guidelines at Ghent University and Ghent University Hospital (UZ-Gent), and were paid for their participation. Written, informed consent was obtained from all participants.

An overview of the three measurement sessions is shown in Table 1, with each session explained in detail in the following sections. Although all participants completed the three sessions, incomplete or corrupt data were found for some participants among the oNH group: subjects 1 and 3 had incomplete AM detection data, subjects 1, 3 and 4 (female) did not perform the SRT processed measurement, and subject 5 (male) was missing WRS data for the HP-filtered speech conditions. The EFR, AM detection, SRT and WRS results in Figures 5-8 are shown without these missing data.

### 7.5 EEG measurements

For the first part of the EEG session, a SAM tone with carrier frequency *f*_*c*_ = 4 kHz, modulation frequency *f*_*m*_ = 120 Hz and *m* = 100% modulation was generated in MATLAB and presented using PLAYREC with a sampling rate of 48 kHz. The processing algorithms were implemented in MATLAB and were applied online to the stimulus before presenting them to the listener. The stimuli were 500-ms long and repeated 1000 times with alternating polarities. A uniformly-distributed, random, silence jitter was applied between consecutive epochs (100 *±* 10 ms) of the 1000 stimulus presentations.

The calibration of the SAM stimuli was carried out based on their peak amplitudes (RMS of the carrier tone), rather than the RMS of each modulated stimulus separately. The reason for avoiding re-calibrating to the RMS was that, after processing the modulated stimulus using our algorithms (Fig. 2), we could keep the same peak amplitude among the different conditions (before and after processing) and thus make a fair comparison between their envelopes and responses. To this end, the *f*_*c*_ = 4 kHz pure-tone carrier that was used to generate the unprocessed SAM tone was calibrated to 70 dB SPL using a B&K sound-level meter type 2606. Then, the calibrated pure tone was modulated to generate the SAM stimulus, which had an RMS of 71.8 dB SPL, and RMS values of 70.3, 70.1 and 68.9 dB SPL after processing with the 13,0,0, 10,0,0 and 7,0,0 conditions, respectively.

For the second part of the session, EEG responses were recorded to the HP-filtered word ‘David’, before and after processing. It has been shown that high-frequency ANFs do not phase lock to the fine-structure content of the sound, but rather follow the envelope changes of the voiced portion of speech [80]. Thus, EFRs to HP-filtered speech stimuli stem from the contribution of mid-frequency fibres and their periodic bursts that follow the fundamental frequency of the stimulus. To this end, the word ‘David’ was selected because of its length and spectral properties, with the voiced parts (/a/ and /i/) containing mostly energy at the fundamental frequency and its harmonics. We isolated the word from a Flemish Matrix sentence [67] and used Praat [105] to monotonise its fundamental frequency *f*_0_ to *∼*220 Hz. Then, the stimulus was HP filtered in the same way as before (*f*_*cut*_ = 1.65 kHz) to remove all content related to the modulation (fundamental) frequency. The sentence was extracted from the calibrated material of the Flemish Matrix, which was based on a 70-dB-SPL calibration of the noise material (SSN) of the test. For the speech stimulus that we used, this resulted in an RMS of 67.9 dB SPL for the unprocessed version and in an RMS of 65.7 dB SPL for the 10,0,0 processed version. The stimuli were 500-ms long (f_s_ = 44,100 Hz) and were repeated 1200 times with alternating polarities. A uniformly-distributed, random, silence jitter was applied between consecutive epochs (200 *±* 20 ms) of the 1200 stimulus presentations.

Scalp-recorded potentials were obtained with a 64-Channel Biosemi EEG recording system and a custom-built trigger-box using a sampling frequency of 16,384 Hz. The electrodes were placed according to the 10-20 standard, using highly conductive gel (Signa gel). The Common-Mode-Sense (CMS) and Driven-Right-Leg DRL) electrodes were placed on top of the head. Two external electrodes were connected to the earlobes as references. During the EFR analysis, all channels were re-referenced to the average of the two earlobe electrodes, with all EFR results representing the Cz channel recordings.

### 7.6 EFR analysis

The EFR analysis was performed according to the procedures of [76]. EEG responses were first filtered with an 800th order Blackman window-based FIR filter. Zero-phase digital filtering was applied between 60 and 600 Hz for the SAM responses, and between 60 and 2500 Hz for the speech responses (to preserve the higher harmonics of the *f*_0_ of the speech stimulus). Signals were broken into epochs of 500 ms relative to the trigger onset, and the first 100 ms of each epoch were omitted in the analysis of the SAM responses. Baseline correction was applied by subtracting the mean of each epoch, before epochs were averaged across trials. Local epoch outliers were then determined using a moving median window of 200 samples (epochs). The peak-to-peak amplitudes of epochs that exceeded the median threshold, corresponding to three times the median absolute deviation, were rejected. An average of 66.4 *±* 36.4 epochs were rejected for the recorded SAM conditions of all participants (6.64 *±* 3.64 %), and an average of 92.4 *±* 53 epochs for the recorded speech conditions (7.7 *±* 4.42 %).

The bootstrapping approach proposed in [106] was employed to estimate the noise-floor (NF) component and remove it from the averaged trials. The bootstrapping procedure was repeated 400 times, resulting in 400 NF-corrected EFR spectra. Each time, n_e_ epochs were drawn randomly with replacement from the epochs after rejection, where n_e_ is the number of remaining epochs after removing the outliers of each measured condition. The FFT spectra were computed with n_FFT_ = 8192 points and the NF component was estimated by repeating the resampling procedure 800 times (with equal numbers of polarities in the draw). A more detailed explanation of the bootstrapping procedure can be found in [59].

After applying the bootstrapping approach, the summed magnitude of the EFR was computed from each of the 400 NF-corrected EFR spectra using:

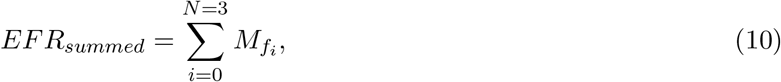

where 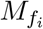 are the EFR spectral-peak (to NF) values at the modulation frequency *f*_0_ of the stimulus and the next *N* = 3 harmonics of *f*_0_. For each bootstrapped EFR spectrum, only the peaks that exceeded one standard deviation of the average NF were considered. Then, the mean and standard deviation across the 400 summed EFR magnitudes was computed to define the mean and variability of the *EFR*_*summed*_ metric (Fig. 5).

### 7.7 AM detection

The AM detection test was implemented as a 3-AFC experiment in the AFC toolbox of MATLAB [104]. Participants were instructed to select the signal that sounded different to the other two, each time choosing between three randomly-presented signals containing two pure tones and a modulated tone of variable modulation depth. An 1-up, 2-down adaptive procedure was used with step sizes of [10, 5, 3, 1] dB and a starting modulation depth of -6 dB. The task was repeated three times for each condition, and the average value and standard deviation of the three trials was calculated (Fig. 6).

In each trial, two 70 dB-SPL, 4-kHz pure tones (reference signals) and a modulated tone (target signal) were generated. The same peak-to-peak calibration as that applied to the EFR stimuli was used, to ensure that the same carrier signal was used among the three signals within each trial. Thus, the modulated tone was calibrated each time to have the same carrier level as the reference pure tones, rather than an RMS calibration that is commonly used in AM detection experiments. As noted above, in the case of a fully-modulated tone calibrated in this way, the difference between the pure-tone and modulated-tone stimuli is less than 2 dB SPL. Even though level cues might be introduced in this worst-case scenario, and during the first trials of the AM detection task, the RMS differences of the modulated stimuli close to the AM detection thresholds are much lower. For the upper limit of our measured AM thresholds (m = 3% / -30.5 dB modulation; Fig. 6), the RMS levels were 70, 69.92, 69.89 and 69.85 dB SPL for the unprocessed, and 13,0,0m, 10,0,0m and 7,0,0m processed stimuli, respectively. Thus, level cues are not expected to have any effect close to the detection thresholds, since the largest difference in this case was *∼*0.15 dB, which is well below the perceivable loudness difference at all frequencies (*∼*0.5 dB [107]).

### 7.8 SI assessment

A custom implementation of the Flemish Matrix Test [67] on the AFC toolbox [104] was used for assessing SI performance. The first test included a standard SRT measurement in BB and HP-filtered speech (before and after processing), while the second test included a word-recognition measurement among the different processing conditions at fixed SNRs of BB and HP-filtered speech. Before presenting them to the listener, the processing algorithms were applied online to the noisy mixture of each sentence. The processing between each step did not significantly increase the total duration of the test. The BB and HP speech material were calibrated based on a separate 70-dB-SPL calibration of the generated BB and HP SSN, respectively, using a B&K sound-level meter type 2606.

At the beginning of each SI test, each subject was presented with a training double list (20 sentences) comprised of all different speech conditions in a random order, so that the subjects could get familiar with the test and the processed speech. For the SRT measurement of each subject, two double lists for BB speech and two double lists for HP speech were randomly presented. For each double list, the SRT measurement was performed using an adaptive procedure to estimate the SNR level that corresponded to *∼*50% word recognition for each subject (SRT). The initial SNR was 0 dB and the noise level was adapted after each trial to reach the desired SNR. For each measured condition, the last 6 reversals were considered to derive the SRT (and deviation) of each participant (Fig. 7), using only the results of the second list for each of the two speech types to account for the training effect of the test [67]. The test was also repeated after applying one processing algorithm (13,0,0m-ref) to the noisy sentences of each trial, with one double list used for each speech type in this case.

During the last SI measurement session, the average WRS was derived for each of the 5 processing conditions (unprocessed, 13,0,0m-ref, 13,0,0, 13,0,0m, 10,0,0m) at 3 SNRs of BB and 3 SNRs of HP-filtered speech. For each participant, the SNRs were individually defined based on the estimated SRT in BB and HP speech (SRT and SRT*±*3 dB). Twelve sentences were randomly presented for each processing condition, resulting in three double lists (5 * 12 = 60 sentences) for each SNR condition that were randomly selected for each run. Then, the average SI score was computed from the 12 measured trials of each condition, separately for each SNR and speech type (Fig. 8).

### 7.9 Statistical analysis

Supplementary Table 4 shows the results of the t-tests that were used to assess the effect of age among the two groups and the effect of processing in each group. To assess the age effect, an independent two-sample t-test was computed between the EFR, AM detection and SRT results of the two groups for each condition (yNH vs. oNH). An independent t-test was not computed between the two groups for the WRS results, since an age-effect comparison is counter-intuitive in this case: The WRS results were measured from the individual SRTs each time, thus offsetting the age-related differences between the groups. For within-group comparisons (processing effect) of the EFRs, AM detection thresholds, SRTs and WRSs, statistically significant differences were assessed using a dependent t-test between each processed condition and the unprocessed result. To account for the multiple comparisons of the unprocessed conditions, Bonferroni correction was applied before establishing statistical significance for the acquired p-values, based on the number of processed conditions that were compared each time. Thus, an *m* value of 3 was used in the cases of SAM EFRs and AM detection thresholds (*m* = 3 hypotheses for each metric). For the WRS results, statistically significant differences were only computed among conditions for which the respective group median increased after processing. This resulted in *m* = 2 comparisons for the WRS results of the yNH group in Fig. 8d, *m* = 3 comparisons for the WRS results of the yNH group in Fig. 8e, and *m* = 1 comparison elsewhere. The *p* values of Supplementary Table 4 were then compared to the corrected significance thresholds (0.05, 0.01, 0.001 and 0.0001 divided by the respective *m* values) to establish statistical significance in Figs. 5-8.

Supplementary Tables 5-9 show the values of Spearman’s rank-order correlation *r* for the SAM EFR, speech EFR, AM detection, SRT and WRS improvement results. Correlations were computed between the yNH and oNH groups separately, between all subjects, and finally between all subjects without taking the four oNH subjects that had HL *>* 20 dB (all w/o oHI) into account. Due to the large number of comparisons, results are reported in each table only for the cases where at least one statistically significant correlation was found. Holm-Bonferroni correction was applied before establishing statistical significance for the acquired p-values, based on the number of comparisons for each metric (*m* = 15 comparisons for each EFR condition, *m* = 18 comparisons for each AM detection condition, *m* = 19 comparisons for each SRT condition and *m* = 25 for the WRS improvement results). The statistical analysis was performed using the Spearman’s *r* implementation from the SciPy Python module, and the p-value correction using the statsmodels Python module. Thus, the correlations with one, two, three and four asterisks in Supplementary Tables 5-9 indicate probability values *p <* 0.05, *p <* 0.01, *p <* 0.001 and *p <* 0.0001, respectively. The number in the parentheses indicate the degrees of freedom (N-2), where N is the number of points used to compute each correlation (sample size). Statistically significant correlations are indicated in bold font. For the EFR conditions, the results of yNH subject 7 were not taken into account in the correlation estimations, since the EFR magnitude was significantly higher than the rest of the subjects for three of the five conditions (cf. Fig. 5).

## 8 Data availability

The evaluation results of our participants can be made available in a public repository upon request or upon acceptance of the paper. Figures 5, 6, 7, 8, 9 and Supplementary Fig. 2 in this paper can be reproduced using this repository.

## 9 Code availability

The MATLAB implementations of the presented hearing-restoration algorithms, as well as the scripts that were used to record and analyse the EFRs, can be made available upon request or upon acceptance of the paper. The source code of the auditory periphery model [50] v1.2 [108] is available via https://doi.org/10.5281/zenodo.3717800 or https://github.com/HearingTechnology/Verhulstetal2018Model.

## 10 Acknowledgments

This work was supported by the European Research Council (ERC) under the Horizon 2020 Research and Innovation Programme (grant agreement No 678120 RobSpear). We thank Heleen Van Der Biest for assisting in the participant recruitment and data collection. English language services were provided by stels-ol.de.

## 11 Author contributions

F.D.: Conceptualisation, Methodology, Software, Validation, Formal analysis, Investigation, Data Curation, Data Collection, Writing: Original Draft, Visualisation; V.V.: Conceptualisation, Methodology, Experimental design; A.O.V.: Software, Experimental design; T.W.: Conceptualisation, Experimental design; S.V.: Conceptualisation, Resources, Supervision, Project administration, Funding acquisition, Writing: Review, Editing.

---

**Supplementary Table 1.** Objective speech evaluation of the different processing strategies. The average PESQ [74] and STOI [73] scores were estimated for the Flemish Matrix sentences [67] of the BB and HP conditions, after adding SSN at SNRs of -8 and -3 dB, respectively (roughly corresponding to the average measured SRTs). In each case, the PESQ and STOI scores were computed using the clean sentence as the reference, with values ranging from 0 (very bad) to 5 (very good) and 0 (very bad) to 1 (very good), respectively.

|  | BB |  | HP |  |
| --- | --- | --- | --- | --- |
|  | PESQ | STOI | PESQ | STOI |
| Unprocessed | 0.6062 | 0.5198 | 1.4806 | 0.7727 |
| 13, 0, 0 | **0.6193** | 0.5085 | 1.3846 | 0.7154 |
| 13, 0, 0m | **0.6145** | 0.5148 | 1.4415 | 0.7454 |
| 13, 0, 0m-ref | **0.6140** | **0.5260** | 1.4421 | 0.7237 |
| 10, 0, 0 | 0.5954 | 0.4637 | 1.2175 | 0.6691 |
| 10, 0, 0m | 0.5731 | 0.4822 | 1.3798 | 0.7298 |
| 10, 0, 0m-ref | 0.5972 | 0.5050 | 1.3509 | 0.7054 |
| 7, 0, 0 | 0.6061 | 0.4469 | 1.0706 | 0.6557 |
| 7, 0, 0m | 0.5105 | 0.4627 | 1.3382 | 0.7294 |
| 7, 0, 0m-ref | 0.5670 | 0.4754 | 1.3062 | 0.7007 |

**Supplementary Table 2.** Fitted parameters of Eqs. *3*-*5* for three CS profiles. The parameters of Eqs. 3-5 were estimated for each CS profile after fitting the functions to the values obtained from the optimisation procedure. The fitted values can be used to compute the parameters *a, b* and *c* for each CS profile.

| CS profile | $a_1$ | $a_2$ | $a_3$ | $a_4$ | $b_1$ | $c_1$ | $c_2$ | $c_3$ | $c_4$ |
| --- | --- | --- | --- | --- | --- | --- | --- | --- | --- |
| 13, 0, 0 | 0.0931 | 2.5804 | 10.2873 | 0.1160 | 7.3543 | -95.8223 | 100.6698 | 283.8136 | 312.0264 |
| 10, 0, 0 | -0.1644 | 0.5473 | 1.5698 | 1.6390 | 39.3868 | -42.7324 | 73.6083 | 13.8984 | 17.5337 |
| 7, 0, 0 | -0.4947 | 0.4224 | 0.6477 | 0.2063 | 291.4200 | 132.0206 | 128.2104 | 20.4308 | 24.6311 |

**Supplementary Table 3.** Parameters for 70 dB-SPL processing. The values of the parameters *b* and *c* of Eq. 2 were computed from Eqs. 4,5 for a 70 dB-SPL SAM tone (*f*_*c*_ = 4 kHz, *f*_*m*_ = 120 Hz). The stimulus envelope had a maximum amplitude of 0.14577. Although the computed value for *a* was close to 1, a constant value of 1 was used across profiles to avoid applying any gain and keep exactly the same peak.

| CS profile | $a$ | $b$ | $c$ |
| --- | --- | --- | --- |
| 13, 0, 0 | 1 | 50.4500 | 4.8476 |
| 10, 0, 0 | 1 | 270.1903 | 30.6769 |
| 7, 0, 0 | 1 | 1999.1160 | 260.1633 |

**Supplementary Figure 1.**
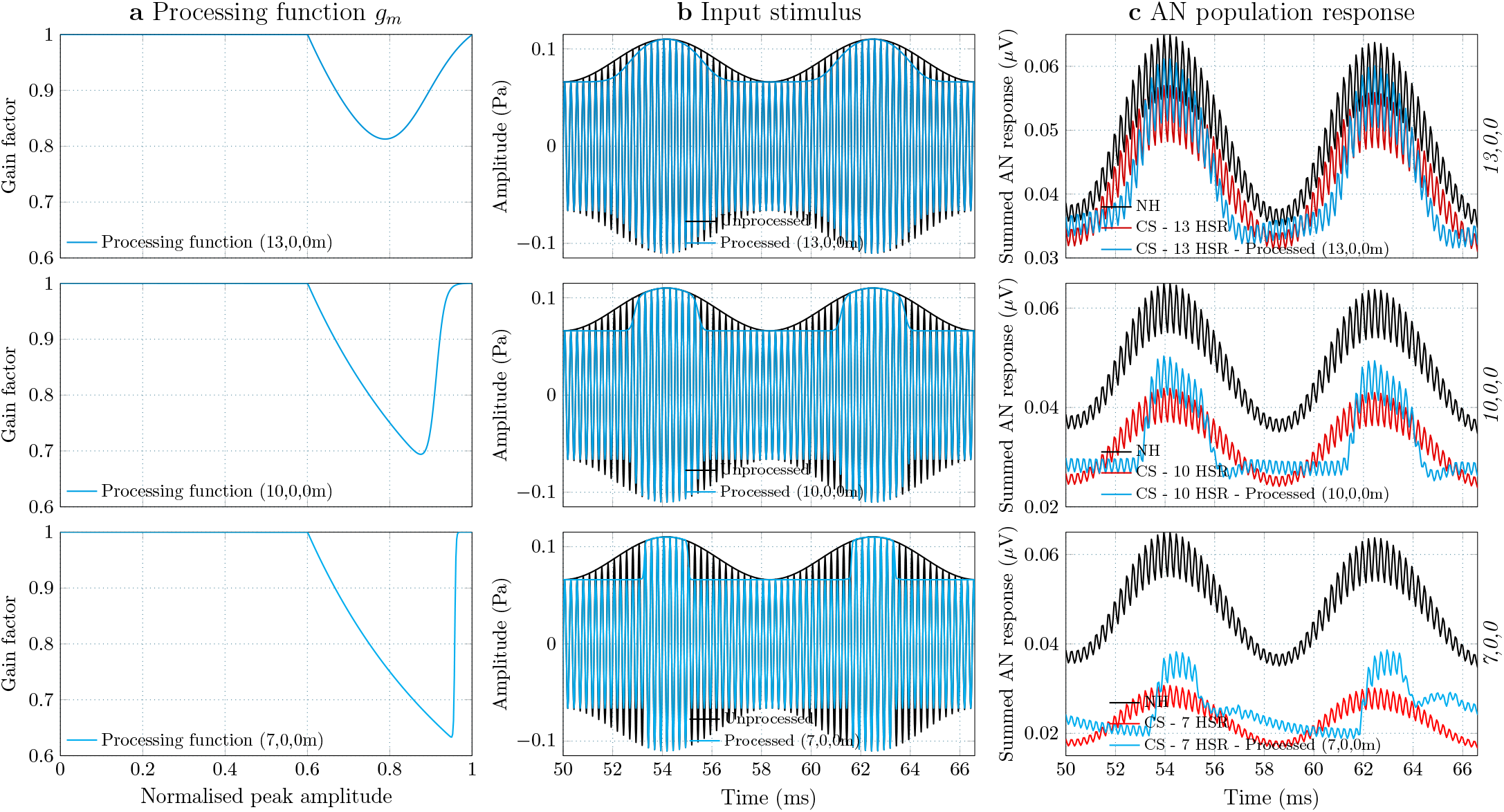
CS compensation of a SAM tone without affecting the modulation depth. The modified processing strategies were applied to a SAM tone stimulus to preserve its modulation depth. The results for a 70-dB-SPL SAM tone with -12 dB modulation depth (*m* = 25 %) are shown. **a** The modified non-linear functions only processed the slope of the envelope, leaving the envelope troughs (and peaks) intact. **b** For each CS type, the non-linear functions were applied to the SAM tones to derive the respective processed stimuli. **c** The simulated AN responses of the NH and CS peripheries are shown in response to the unprocessed SAM stimulus and the respective processed version, given as input to the corresponding CS periphery.

**Supplementary Figure 2.**
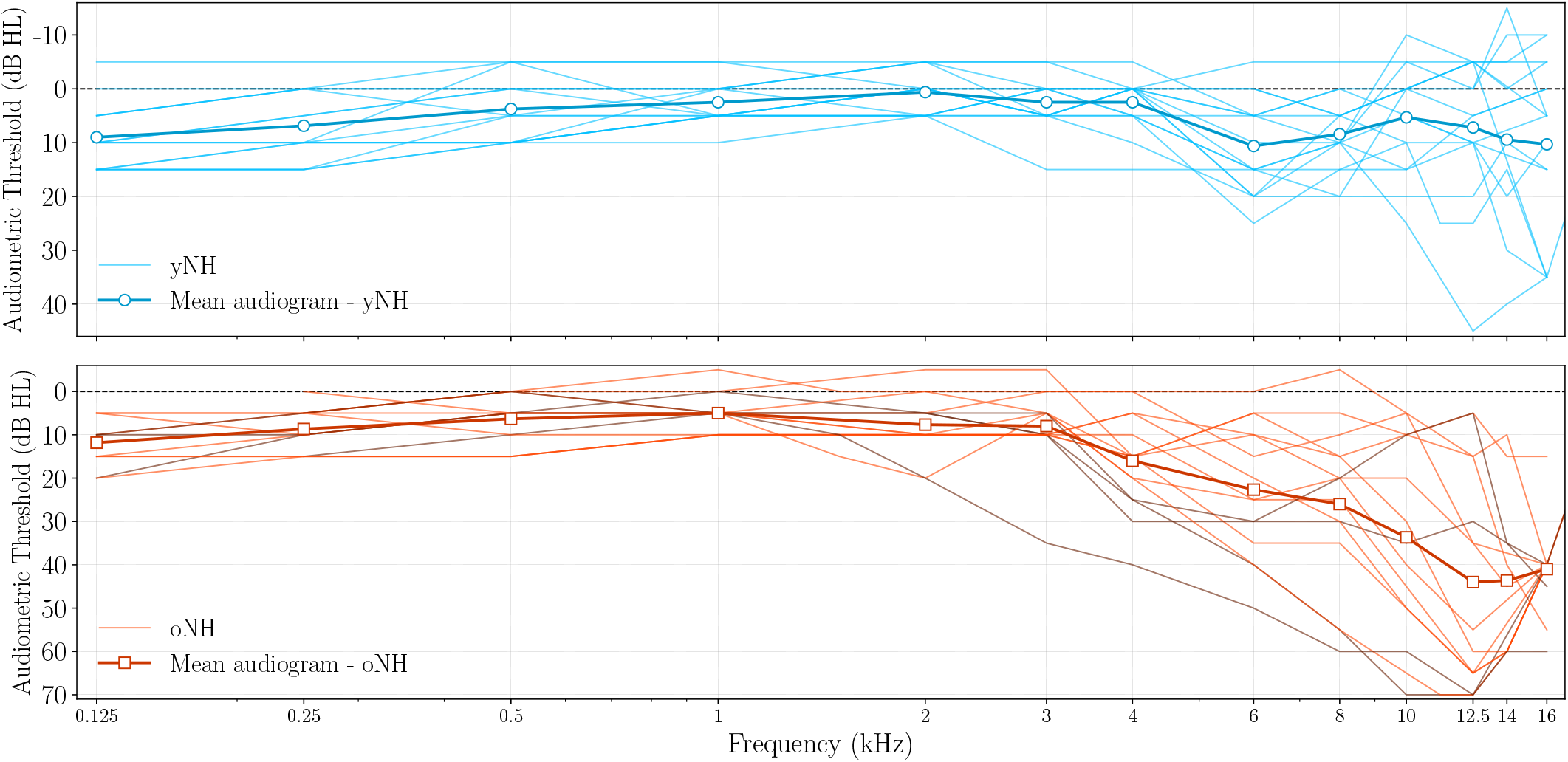
Audiograms of participants. The individual audiograms of the participants are shown in thin lines for the two age groups. The thick lines represent the grand-averaged audiogram for each age group. The darker thin lines for the oNH group indicate participants with HL *>* 20 dB at frequencies up to 4 kHz.

**Supplementary Figure 3.**
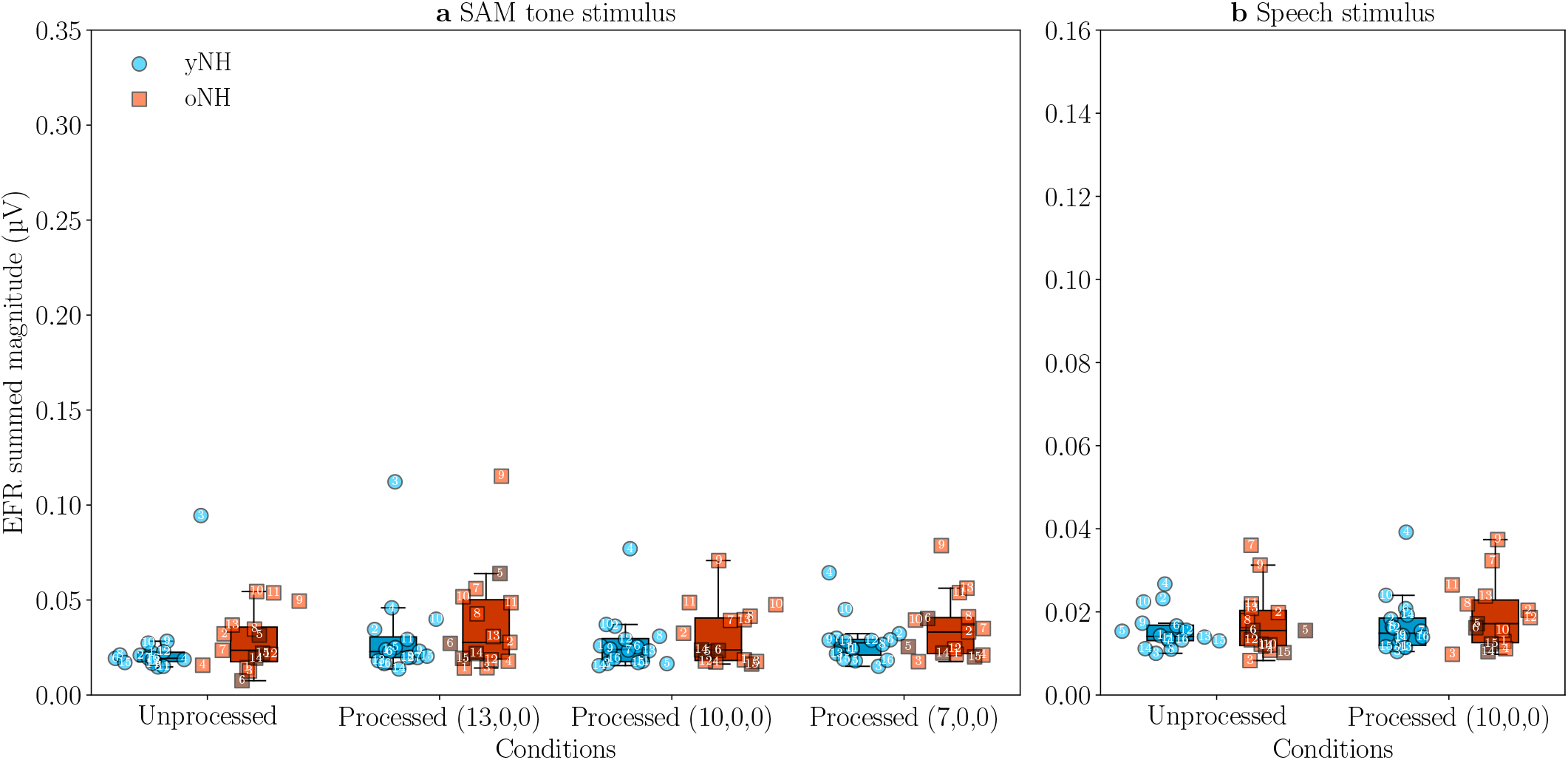
Estimated EFR noise-floor (NF) magnitudes (cf. Fig. 5). The magnitudes of the estimated EFR NFs are shown for the SAM and speech stimuli before and after processing, and were subtracted from the recorded EFR results to yield the EFR magnitudes of Fig. 5 (see Methods). In both panels, the individual points correspond to the mean and standard deviation of the summed EFR magnitudes (EFR_summed_), computed from the EFR noise spectra of each participant (Eq. 10). The box and whisker plots indicate the EFR medians and quartiles of the two groups under each condition.

**Supplementary Table 4.** T-test results. Computed *t* statistics and *p* values for the measured results of the two age groups. An independent two-sample t-test was used to assess statistical significance between the group results of each condition (age effect). A dependent t-test was used to assess statistical significance after processing among the different conditions of each group (processing effect), each time computed between the respective processed result and the unprocessed reference. In the case of the WRS results, an independent t-test was not used, while the processing effect was assessed only for the cases where a higher score median was obtained after processing.

| Condition | Age effect (yNH vs. oNH) | Processing effect (yNH) | Processing effect (oNH) |
| --- | --- | --- | --- |
| EFR SAM (unprocessed) | $t_{(14)}=2.8367, p=0.0083$ | - | - |
| EFR SAM (13,0,0) | $t_{(14)}=1.5130, p=0.1412$ | $t_{(14)}=-0.4882, p=0.6325$ | $t_{(13)}=-1.9284, p=0.0743$ |
| EFR SAM (10,0,0) | $t_{(14)}=2.2441, p=0.0349$ | $t_{(14)}=-5.0698, p=0.0001$ | $t_{(13)}=-2.3656, p=0.0330$ |
| EFR SAM (7,0,0) | $t_{(14)}=3.4871, p=0.0023$ | $t_{(14)}=-5.1887, p=0.0001$ | $t_{(13)}=-5.6619, p=0.0001$ |
| EFR speech (unprocessed) | $t_{(14)}=4.2434, p=0.0005$ | - | - |
| EFR speech (10,0,0) | $t_{(14)}=3.0411, p=0.0069$ | $t_{(14)}=-3.1010, p=0.0073$ | $t_{(13)}=-4.4866, p=0.0005$ |
| AM thresholds (unprocessed) | $t_{(14)}=-1.3291, p=0.1957$ | - | - |
| AM thresholds (13,0,0) | $t_{(14)}=-1.8179, p=0.0806$ | $t_{(14)}=5.4243, p=0.0001$ | $t_{(11)}=2.3224, p=0.0386$ |
| AM thresholds (10,0,0) | $t_{(14)}=-2.8228, p=0.0090$ | $t_{(14)}=5.5684, p=0.0001$ | $t_{(11)}=4.9300, p=0.0003$ |
| AM thresholds (7,0,0) | $t_{(14)}=-2.8346, p=0.0087$ | $t_{(14)}=9.2074, p=0.0000$ | $t_{(11)}=7.9724, p=0.0000$ |
| SRT BB (unprocessed) | $t_{(14)}=-1.1889, p=0.2450$ | - | - |
| SRT BB (13,0,0m-ref) | $t_{(14)}=-1.6882, p=0.1042$ | $t_{(14)}=1.4250, p=0.1746$ | $t_{(10)}=-0.5880, p=0.5684$ |
| SRT HP (unprocessed) | $t_{(14)}=-2.3250, p=0.0274$ | - | - |
| SRT HP (13,0,0m-ref) | $t_{(14)}=-2.0321, p=0.0525$ | $t_{(14)}=-1.0796, p=0.2974$ | $t_{(10)}=-2.2606, p=0.0450$ |
| WRS (13,0,0m) - BB (SRT-3) | - | $t_{(14)}=-2.7296, p=0.0155$ | $t_{(13)}=-1.8708, p=0.0824$ |
| WRS (13,0,0m) - BB (SRT) | - | $t_{(14)}=-2.1877, p=0.0449$ | $t_{(13)}=1.7425, p=0.1033$ |
| WRS (13,0,0m-ref) - BB (SRT+3) | - | - | $t_{(13)}=0.5214, p=0.6102$ |
| WRS (13,0,0) - HP (SRT-3) | - | $t_{(14)}=-0.9802, p=0.3425$ | - |
| WRS (13,0,0m) - HP (SRT-3) | - | $t_{(14)}=-3.5615, p=0.0028$ | $t_{(12)}=-0.6740, p=0.5121$ |
| WRS (13,0,0) - HP (SRT) | - | $t_{(14)}=-0.6115, p=0.5500$ | - |
| WRS (13,0,0m) - HP (SRT) | - | $t_{(14)}=-0.8686, p=0.3987$ | - |
| WRS (10,0,0m) - HP (SRT) | - | $t_{(14)}=0.0000, p=1.0000$ | - |
| WRS (13,0,0m) - HP (SRT+3) | - | $t_{(14)}=-1.3547, p=0.1956$ | - |

**Supplementary Table 5.**
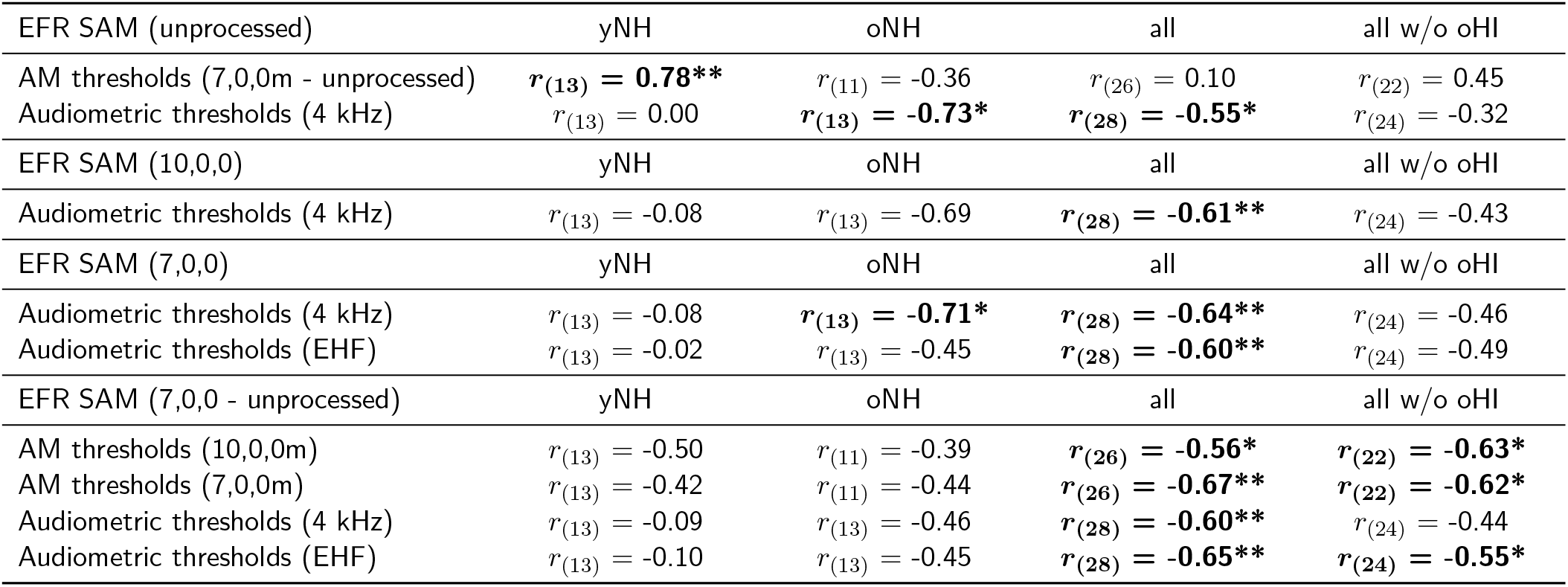
Significant correlations of SAM EFRs. Spearman’s *r* values between the EFR results of the four SAM conditions and the measured AM detection thresholds, SRTs and audiometric thresholds. The table shows only the cases for which at least one statistically significant correlation was found (in bold), with the asterisks corresponding to the significance after the Holm-Bonferroni correction.

| EFR SAM (unprocessed) | yNH | oNH | all | all w/o oHI |
| --- | --- | --- | --- | --- |
| AM thresholds (7,0,0m - unprocessed) | **$r_{(13)} = 0.78^{**}$** | $r_{(11)} = -0.36$ | $r_{(26)} = 0.10$ | $r_{(22)} = 0.45$ |
| Audiometric thresholds (4 kHz) | $r_{(13)} = 0.00$ | **$r_{(13)} = -0.73^*$** | **$r_{(28)} = -0.55^*$** | $r_{(24)} = -0.32$ |
| EFR SAM (10,0,0) | yNH | oNH | all | all w/o oHI |
| Audiometric thresholds (4 kHz) | $r_{(13)} = -0.08$ | $r_{(13)} = -0.69$ | **$r_{(28)} = -0.61^{**}$** | $r_{(24)} = -0.43$ |
| EFR SAM (7,0,0) | yNH | oNH | all | all w/o oHI |
| Audiometric thresholds (4 kHz) | $r_{(13)} = -0.08$ | **$r_{(13)} = -0.71^*$** | **$r_{(28)} = -0.64^{**}$** | $r_{(24)} = -0.46$ |
| Audiometric thresholds (EHF) | $r_{(13)} = -0.02$ | $r_{(13)} = -0.45$ | **$r_{(28)} = -0.60^{**}$** | $r_{(24)} = -0.49$ |
| EFR SAM (7,0,0 - unprocessed) | yNH | oNH | all | all w/o oHI |
| AM thresholds (10,0,0m) | $r_{(13)} = -0.50$ | $r_{(11)} = -0.39$ | **$r_{(26)} = -0.56^*$** | **$r_{(22)} = -0.63^*$** |
| AM thresholds (7,0,0m) | $r_{(13)} = -0.42$ | $r_{(11)} = -0.44$ | **$r_{(26)} = -0.67^{**}$** | **$r_{(22)} = -0.62^*$** |
| Audiometric thresholds (4 kHz) | $r_{(13)} = -0.09$ | $r_{(13)} = -0.46$ | **$r_{(28)} = -0.60^{**}$** | $r_{(24)} = -0.44$ |
| Audiometric thresholds (EHF) | $r_{(13)} = -0.10$ | $r_{(13)} = -0.45$ | **$r_{(28)} = -0.65^{**}$** | **$r_{(24)} = -0.55^*$** |

**Supplementary Table 6.** Significant correlations of speech EFRs. Spearman’s *r* values between the EFR results of the two speech conditions and the measured AM detection thresholds, SRTs and audiometric thresholds. The table shows only the cases for which at least one statistically significant correlation was found (in bold), with the asterisks corresponding to the significance after the Holm-Bonferroni correction.

| EFR speech (unprocessed) | yNH | oNH | all | all w/o oHI |
| --- | --- | --- | --- | --- |
| AM thresholds (10,0,0m - unprocessed) | $r_{(13)} = 0.30$ | $r_{(11)} = \mathbf{-0.75^*}$ | $r_{(26)} = -0.21$ | $r_{(22)} = 0.05$ |
| AM thresholds (7,0,0m - unprocessed) | $r_{(13)} = 0.44$ | $r_{(11)} = \mathbf{-0.86^{**}}$ | $r_{(26)} = -0.23$ | $r_{(22)} = 0.03$ |
| Audiometric thresholds (4 kHz) | $r_{(13)} = -0.10$ | $r_{(13)} = -0.67$ | $r_{(28)} = \mathbf{-0.69^{***}}$ | $r_{(24)} = -0.54$ |
| Audiometric thresholds (EHF) | $r_{(13)} = 0.05$ | $r_{(13)} = -0.58$ | $r_{(28)} = \mathbf{-0.68^{***}}$ | $r_{(24)} = \mathbf{-0.60^*}$ |
| EFR speech (10,0,0) | yNH | oNH | all | all w/o oHI |
| AM thresholds (10,0,0m - unprocessed) | $r_{(13)} = 0.30$ | $r_{(11)} = \mathbf{-0.81^*}$ | $r_{(26)} = -0.25$ | $r_{(22)} = -0.00$ |
| AM thresholds (7,0,0m - unprocessed) | $r_{(13)} = 0.39$ | $r_{(11)} = \mathbf{-0.79^*}$ | $r_{(26)} = -0.21$ | $r_{(22)} = 0.06$ |
| Audiometric thresholds (4 kHz) | $r_{(13)} = 0.05$ | $r_{(13)} = \mathbf{-0.70^*}$ | $r_{(28)} = \mathbf{-0.63^{**}}$ | $r_{(24)} = -0.44$ |

**Supplementary Table 7.** Significant correlations of AM detection thresholds. Spearman’s *r* values between the AM detection thresholds of the four conditions and the measured EFRs, SRTs and audiometric thresholds. The table shows only the cases for which at least one statistically significant correlation was found (in bold), with the asterisks corresponding to the significance after the Holm-Bonferroni correction.

| AM thresholds (13,0,0m - unprocessed) | yNH | oNH | all | all w/o oHI |
| --- | --- | --- | --- | --- |
| SRT HP (13,0,0m-ref) | $r_{(14)} = 0.63$ | $r_{(10)} = 0.48$ | $r_{(26)} = \mathbf{0.64^{**}}$ | $r_{(22)} = \mathbf{0.60^*}$ |
| SRT HP (13,0,0m-ref - unprocessed) | $r_{(14)} = 0.60$ | $r_{(10)} = 0.43$ | $r_{(26)} = \mathbf{0.58^*}$ | $r_{(22)} = 0.51$ |
| AM thresholds (10,0,0m) | yNH | oNH | all | all w/o oHI |
| EFR SAM (7,0,0 - unprocessed) | $r_{(13)} = -0.50$ | $r_{(11)} = -0.39$ | $r_{(26)} = \mathbf{-0.56^*}$ | $r_{(22)} = \mathbf{-0.63^*}$ |
| SRT BB (13,0,0m-ref) | $r_{(14)} = \mathbf{-0.80^{**}}$ | $r_{(10)} = 0.25$ | $r_{(26)} = -0.09$ | $r_{(22)} = -0.29$ |
| SRT HP (unprocessed) | $r_{(14)} = \mathbf{-0.81^{**}}$ | $r_{(11)} = -0.15$ | $r_{(27)} = -0.14$ | $r_{(23)} = -0.23$ |
| SRT HP (13,0,0m-ref) | $r_{(14)} = \mathbf{-0.75^*}$ | $r_{(10)} = 0.25$ | $r_{(26)} = -0.17$ | $r_{(22)} = -0.36$ |
| SRT BB (13,0,0m-ref - unprocessed) | $r_{(14)} = \mathbf{-0.70^*}$ | $r_{(10)} = -0.23$ | $r_{(26)} = -0.33$ | $r_{(22)} = -0.25$ |
| AM thresholds (10,0,0m - unprocessed) | yNH | oNH | all | all w/o oHI |
| EFR speech (10,0,0) | $r_{(13)} = 0.30$ | $r_{(11)} = \mathbf{-0.81^*}$ | $r_{(26)} = -0.25$ | $r_{(22)} = -0.00$ |
| Audiometric thresholds (EHF) | $r_{(14)} = 0.04$ | $r_{(11)} = \mathbf{0.77^*}$ | $r_{(27)} = 0.30$ | $r_{(23)} = 0.11$ |
| AM thresholds (7,0,0m) | yNH | oNH | all | all w/o oHI |
| EFR SAM (7,0,0 - unprocessed) | $r_{(13)} = -0.42$ | $r_{(11)} = -0.44$ | $r_{(26)} = \mathbf{-0.67^{**}}$ | $r_{(22)} = \mathbf{-0.62^*}$ |
| Audiometric thresholds (4 kHz) | $r_{(14)} = 0.40$ | $r_{(11)} = 0.15$ | $r_{(27)} = \mathbf{0.59^*}$ | $r_{(23)} = 0.56$ |
| Audiometric thresholds (EHF) | $r_{(14)} = 0.18$ | $r_{(11)} = 0.46$ | $r_{(27)} = \mathbf{0.68^{***}}$ | $r_{(23)} = \mathbf{0.62^*}$ |
| AM thresholds (7,0,0m - unprocessed) | yNH | oNH | all | all w/o oHI |
| EFR SAM (unprocessed) | $r_{(13)} = \mathbf{0.78^*}$ | $r_{(11)} = -0.36$ | $r_{(26)} = 0.10$ | $r_{(22)} = 0.45$ |
| EFR speech (unprocessed) | $r_{(13)} = 0.44$ | $r_{(11)} = \mathbf{-0.86^{**}}$ | $r_{(26)} = -0.23$ | $r_{(22)} = 0.03$ |
| EFR speech (10,0,0) | $r_{(13)} = 0.39$ | $r_{(11)} = \mathbf{-0.79^*}$ | $r_{(26)} = -0.21$ | $r_{(22)} = 0.06$ |

**Supplementary Table 8.** Significant correlations of SRTs. Spearman’s *r* values between the SRTs and the measured EFRs, AM detection thresholds and audiometric thresholds. The table shows only the cases for which at least one statistically significant correlation was found (in bold), with the asterisks corresponding to the significance after the Holm-Bonferroni correction.

| SRT BB (13,0,0m-ref) | yNH | oNH | all | all w/o oHI |
| --- | --- | --- | --- | --- |
| AM thresholds (10,0,0m) | $r_{(14)} = \mathbf{-0.80^{**}}$ | $r_{(10)} = 0.25$ | $r_{(26)} = -0.09$ | $r_{(22)} = -0.29$ |
| SRT BB (13,0,0m-ref - unprocessed) | yNH | oNH | all | all w/o oHI |
| AM thresholds (10,0,0m) | $r_{(14)} = \mathbf{-0.70^*}$ | $r_{(10)} = -0.23$ | $r_{(26)} = -0.33$ | $r_{(22)} = -0.25$ |
| SRT HP (unprocessed) | yNH | oNH | all | all w/o oHI |
| AM thresholds (10,0,0m) | $r_{(14)} = \mathbf{-0.81^{**}}$ | $r_{(11)} = -0.15$ | $r_{(27)} = -0.14$ | $r_{(23)} = -0.23$ |
| SRT HP (13,0,0m-ref) | yNH | oNH | all | all w/o oHI |
| AM thresholds (10,0,0m) | $r_{(14)} = \mathbf{-0.75^*}$ | $r_{(10)} = 0.25$ | $r_{(26)} = -0.17$ | $r_{(22)} = -0.36$ |
| AM thresholds (13,0,0m - unprocessed) | $r_{(14)} = 0.63$ | $r_{(10)} = 0.48$ | $r_{(26)} = \mathbf{0.64^{**}}$ | $r_{(22)} = \mathbf{0.60^*}$ |
| SRT HP (13,0,0m-ref - unprocessed) | yNH | oNH | all | all w/o oHI |
| AM thresholds (13,0,0m - unprocessed) | $r_{(14)} = 0.60$ | $r_{(10)} = 0.43$ | $r_{(26)} = \mathbf{0.58^*}$ | $r_{(22)} = 0.51$ |

**Supplementary Table 9.** Significant correlations of the WRS improvement in the six SNR conditions. Spearman’s *r* values between the WRS improvement of the processing algorithms and the measured EFRs, AM detection thresholds, SRTs and audiometric thresholds. The table shows only the cases for which at least one statistically significant correlation was found (in bold), with the asterisks corresponding to the significance after the Holm-Bonferroni correction.

| BB (SRT-3 dB) |  |  |  |  |
| --- | --- | --- | --- | --- |
| WRS (13,0,0m-ref - unprocessed) | yNH | oNH | all | all w/o oHI |
| AM thresholds (7,0,0m) | $r_{(14)} = \mathbf{-0.75^*}$ | $r_{(11)} = 0.39$ | $r_{(27)} = -0.21$ | $r_{(23)} = -0.20$ |
| HP (SRT-3 dB) |  |  |  |  |
| WRS (13,0,0m-ref - unprocessed) | yNH | oNH | all | all w/o oHI |
| AM thresholds (7,0,0m) | $r_{(14)} = -0.59$ | $r_{(10)} = -0.25$ | $r_{(26)} = \mathbf{-0.57^*}$ | $r_{(23)} = \mathbf{-0.59^*}$ |
| Audiometric thresholds (4 kHz) | $r_{(14)} = -0.57$ | $r_{(12)} = -0.59$ | $r_{(28)} = \mathbf{-0.54^*}$ | $r_{(25)} = \mathbf{-0.61^*}$ |
| WRS (13,0,0 - unprocessed) | yNH | oNH | all | all w/o oHI |
| EFR SAM (unprocessed) | $r_{(13)} = \mathbf{-0.85^{**}}$ | $r_{(12)} = -0.25$ | $r_{(27)} = -0.34$ | $r_{(24)} = -0.38$ |
| WRS (13,0,0m - unprocessed) | yNH | oNH | all | all w/o oHI |
| EFR speech (unprocessed) | $r_{(13)} = \mathbf{-0.84^{**}}$ | $r_{(12)} = -0.08$ | $r_{(27)} = -0.13$ | $r_{(24)} = -0.11$ |
| HP (SRT+3 dB) |  |  |  |  |
| WRS (13,0,0m-ref - unprocessed) | yNH | oNH | all | all w/o oHI |
| EFR SAM (unprocessed) | $r_{(13)} = \mathbf{-0.75^*}$ | $r_{(12)} = -0.50$ | $r_{(27)} = -0.47$ | $r_{(24)} = -0.46$ |

**Supplementary Table 10.** Word-recognition improvement for the 13,0,0m strategy (cf. Fig. 8). The median SI improvement achieved with the 13,0,0m strategy is shown for each SNR condition, after subtracting the median WRS of the unprocessed condition. The median scores were defined from the average results of all subjects in each group, which were computed either using all 12 trials of each subject (trials 1-12) or after omitting the first 4 trials of the results (trials 5-12). For each group in each condition, the best SI improvement is indicated by bold font.

|  | BB |  |  | HP |  |  |
| --- | --- | --- | --- | --- | --- | --- |
|  | SRT -3 dB | SRT | SRT +3 dB | SRT -3 dB | SRT | SRT +3 dB |
| yNH (trials 1-12) | 5 % | **0.83 %** | 0 % | 8.33 % | 4.17 % | **3.33 %** |
| oNH (trials 1-12) | 3.33 % | **1.67 %** | 0 % | 3.33 % | -1.67 % | -3.33 % |
| yNH (trials 5-12) | **6.25 %** | 0 % | **1.25 %** | **10 %** | **7.5 %** | 0 % |
| oNH (trials 5-12) | **7.5 %** | -2.5 % | **2.5 %** | **3.75 %** | **2.5 %** | **-1.25 %** |

